# Immune-metabolic-redox ecosystems define spatially organized tumor states in head and neck squamous cell carcinoma

**DOI:** 10.64898/2026.08.31.746545

**Authors:** Kirtikar Shukla

**Affiliations:** Department of Internal Medicine, Section on Molecular Medicine, Wake Forest University School of Medicine, Winston-Salem, North Carolina, USA

**Keywords:** Head and neck squamous cell carcinoma (HNSCC), Spatial transcriptomics, Tumor microenvironment, Immune ecosystems, Immunometabolism, Oxidative stress signaling, Tumor-immune interactions, Single-cell spatial analysis, Immune-Metabolic-Redox Ecosystem Score (IMRES), Immune heterogeneity

## Abstract

**Background:** Spatial organization is increasingly recognized as a key determinant of tumor-immune interactions in head and neck squamous cell carcinoma (HNSCC). The GSE300147 Xenium spatial transcriptomic resource generated by McCord and colleagues established a framework for mapping spatially coordinated T-cell states in HNSCC. However, how tumor-enriched epithelial immune states relate to metabolic, redox, and stress-adaptive transcript programs remains incompletely defined.

**Methods:** A secondary, data-driven reanalysis of GSE300147 was performed, focusing on 17 confirmed HNSCC Xenium sections after exclusion of a non-HNSCC ameloblastoma specimen. A total of 1,148,244 cells were analyzed, including 558,867 EpCAM^+^ tumor-enriched epithelial cells. Tumor-enriched epithelial cells were classified into Hot, Intermediate, and Cold states using a Composite Hotness framework integrating T-cell inflammatory signature score, checkpoint-associated signaling, CD274 expression, IFN/antigen-presentation signature score (IFN/AP), and tumor-immune proximity. Six metabolic ecosystem states, neighborhood profiling, spatial permutation testing, and an integrated Immune-Metabolic-Redox Ecosystem Score (IMRES) were then applied.

**Results:** Immune activation was spatially heterogeneous across HNSCC sections. Immune-hot tumor-enriched epithelial regions showed not only inflammatory, checkpoint-associated, and antigen-presentation signature scores, but also coordinated metabolic, oxidative-redox, and stress-response transcript programs. IMRES, derived from available immune, metabolic, redox, and stress-response transcript components represented in the Xenium panel, increased progressively from Cold to Intermediate to Hot tumor-enriched epithelial states and was associated with NFE2L2, GDF15, HLA-DRA, CD274, KEAP1, and MDM2. Integrating IMRES with Composite Hotness identified a distinct Hot+IMRES_high_ ecosystem comprising 106,874 tumor-enriched epithelial cells. This state showed the strongest immune-active and stress-adaptive features and was positioned closer to immune populations than expected by random assignment. An alternative rank-based robustness analysis reproduced the IMRES-associated ecosystem axis and correlated with the original module-based score (Spearman r = 0.597).

**Conclusions:** This secondary reanalysis extends the original spatial T-cell framework by defining a complementary tumor-centered immune-metabolic-redox ecosystem in HNSCC. IMRES provides a transcript-derived framework for identifying Hot+IMRES_high_ neighborhoods where immune activation, checkpoint signaling, metabolic remodeling, and stress adaptation converge, providing a hypothesis-generating framework for studying immune resistance and therapeutic vulnerability.

## 1. Introduction

HNSCC remains difficult to treat because of marked tumor heterogeneity and a complex tumor microenvironment. Although multimodal therapy and PD-1 inhibitors such as pembrolizumab and nivolumab have improved outcomes for some patients, durable responses remain limited, with objective response rates often around 14%–22%(1, 2). Available biomarkers, including PD-L1 expression, tumor mutational burden, and measures of immune-cell infiltration, provide useful information but do not fully capture the local tissue environments that influence antitumor immunity(3–5). Increasing evidence suggests that the spatial organization of immune and tumor cells within the tumor microenvironment plays an important role in determining immune function, therapeutic sensitivity, and disease progression(6, 7). Spatial transcriptomic technologies have provided new opportunities to investigate these local tissue environments while preserving cellular architecture(8, 9). Unlike bulk RNA sequencing or dissociated single-cell approaches, spatial methods retain positional information and allow direct assessment of tumor–immune interactions within intact tissue sections(8, 10). Recent studies in HNSCC and other solid tumors have shown that the spatial arrangement of immune cells, stromal populations, and tumor-enriched epithelial compartments can influence immune activation, T-cell exhaustion, and response to therapy(11, 12). These observations support the view that antitumor immunity is shaped not only by the presence of immune cells but also by their spatial relationships with surrounding tumor ecosystems(8, 12). In parallel with advances in spatial biology, HNSCC has increasingly been recognized as a metabolically adaptive malignancy(13). Alterations in lipid metabolism, cholesterol homeostasis, prostaglandin and eicosanoid signaling, oxidative stress responses, and mitochondrial function have all been linked to tumor progression and treatment resistance(14, 15). Several studies have demonstrated that NRF2 signaling and associated antioxidant pathways promote adaptation to oxidative stress, whereas ferroptosis-associated pathways influence tumor-cell survival and therapeutic responsiveness(16, 17). In addition, mitochondrial stress responses and radiation-associated stress programs have emerged as important regulators of tumor adaptation to environmental and therapeutic pressures(17, 18). Although these pathways have been studied extensively in bulk tumors, cell lines, and conventional single-cell datasets, their spatial relationship to immune-active tumor regions remains poorly understood(19, 20). In particular, it remains unclear whether immune-inflamed tumor neighborhoods coexist with metabolic and stress-adaptive programs that could simultaneously support immune engagement and therapeutic resistance(21, 22). The public GSE300147 resource generated by McCord and colleagues offers a unique opportunity to address this question(23). Using Xenium spatial transcriptomics combined with patient-specific T-cell receptor probes, the original study established a framework for mapping clonally related T cells within HNSCC tissue and demonstrated that T-cell transcriptional states vary according to spatial location(23, 24). Presumed tumor-reactive T cells were broadly distributed throughout the tumor microenvironment, and cells belonging to the same T-cell clone occupied distinct functional states depending on their local surroundings(25, 26). These findings highlighted the importance of spatial context in shaping antitumor immune responses. The present study builds on that spatial framework but asks a different biological question. Rather than focusing on T-cell clonality, this reanalysis examined how tumor-enriched epithelial cells are organized across immune-hot and immune-cold spatial states and whether these states are associated with coordinated metabolic, redox, and stress-response programs(27–29). Using tumor-centered analyses of 558,867 EpCAM^+^ tumor-enriched epithelial cells from 17 HNSCC Xenium sections, immune activation, checkpoint signaling, tumor–immune proximity, lipid and cholesterol metabolism, oxidative-redox adaptation, ferroptosis-associated programs, and radiation-stress signatures were integrated into a unified Immune-Metabolic-Redox Ecosystem Score (IMRES), while additional stress-response programs, including mitochondrial-stress signatures, were evaluated in parallel(30–33). Spatial neighborhood analysis, ecosystem-state modeling, permutation-based spatial validation, and rank-based robustness testing were then used to evaluate whether immune-hot tumor-enriched epithelial regions represent distinct biological ecosystems rather than isolated inflammatory cell states(34, 35). By combining spatial immune-state classification with metabolic and stress-response profiling, this study extends the value of GSE300147 beyond T-cell mapping and provides a complementary perspective on how immune activation, metabolic remodeling, and stress adaptation are organized within the HNSCC tumor microenvironment. The findings support a model in which immune-hot tumor-enriched epithelial regions are embedded within spatially coordinated immune-metabolic-redox ecosystems that may be relevant to antitumor immunity and mechanisms of therapeutic resistance.

## 2. Materials and Methods

### 2.1 Data source and study design

This study was performed as a secondary computational reanalysis of the publicly available Xenium spatial transcriptomic dataset GSE300147, deposited in the NCBI Gene Expression Omnibus(23). The dataset was originally generated by McCord et al(23) for the study Single-cell TCR mapping reveals spatially coordinated T cell states in head and neck cancer, published in Science Immunology. The original study introduced a spatial T-cell receptor profiling strategy that linked clonal identity, transcriptional state, and spatial position of individual T cells in the HNSCC tumor microenvironment. The GEO record for GSE300147 describes Xenium spatial transcriptomic profiling of ten HNSCC patient tumors and one ameloblastoma tumor. Seven of the ten HNSCC tumors were profiled as replicate tissue sections on different slides and days, resulting in 17 HNSCC Xenium sections and one ameloblastoma section. The dataset used the 10x Genomics human multi-tissue and cancer gene expression panel together with custom probes targeting patient-specific TCR CDR3 regions, additional T-cell genes, HPV oncoprotein genes, and tumor genes of interest. The present study did not generate the original patient cohort, tissue sections, Xenium assays, TCR probes, flow cytometry, single-cell RNA sequencing, or TCR sequencing data. These experimental components were part of the original McCord et al. resource(23). The current analysis focused on a tumor-centered secondary reanalysis of the processed Xenium transcriptomic data, with emphasis on immune-hot and immune-cold tumor ecosystems, metabolic and redox transcript programs, tumor-immune spatial proximity, and integrated immune-metabolic-redox scoring. Because the aim of this study was to interpret HNSCC biology, the ameloblastoma specimen, corresponding to GSM9054488, was excluded from all downstream analyses(23). The final analysis therefore included 17 HNSCC spatial sections. Across these sections, the processed analysis contained 1,148,244 total cells, of which 558,867 EpCAM^+^ tumor-enriched epithelial cells were retained for the tumor-centered analyses.

### 2.2 Computational environment and data handling

Processed Xenium-derived AnnData objects were used as the starting point for all analyses. These files contained transcript count matrices, cell-level metadata, segmentation-derived information, sample identifiers, and spatial coordinates. Analyses were performed in Python 3.9 using Scanpy, AnnData, NumPy, Pandas, SciPy, scikit-learn, Matplotlib, and Seaborn (36, 37). Scanpy was used to handle the large single-cell expression matrices and associated metadata structures(36). The Xenium platform is an imaging-based spatial transcriptomic approach that detects targeted transcripts in situ and assigns RNA molecules to their spatial location within tissue sections(9, 38). In this study, Xenium measurements were interpreted as targeted transcript-level spatial readouts, not as direct measurements of protein abundance, metabolite concentrations, lipid mediators, or functional pathway activity(38).

### 2.3 Identification of tumor-enriched epithelial cells

Tumor-centered analyses were restricted to epithelial-enriched compartments. Cells with detectable EpCAM transcript expression were classified as tumor-enriched epithelial cells and retained for downstream analyses(20). This filtering strategy was used to enrich the analytical population for epithelial tumor-associated compartments within the Xenium data. EpCAM^+^ was not treated as definitive malignant-cell calling(39, 40). No copy-number inference, mutation-based tumor-cell classification, lineage tracing, or histopathological reannotation was performed in this secondary analysis. For this reason, these cells are referred to throughout the manuscript as tumor-enriched epithelial cells rather than definitively malignant cells.

### 2.4 Composite Hotness Score and classification of immune ecosystem states

A Composite Hotness Score was developed to quantify local immune activation within tumor-enriched epithelial cells, building on spatial tumor-immune analysis frameworks and the original GSE300147 HNSCC Xenium resource(6–8, 12, 23, 28, 34). The score integrated five features: T-cell inflammatory signature score, checkpoint-associated signaling, CD274 expression, IFN/antigen-presentation signature score, and tumor-immune spatial proximity. Each feature was standardized before integration into a single composite score. Tumor-enriched epithelial cells were ranked according to Composite Hotness Score(41, 42). Cells in the upper quartile were classified as Hot, cells in the lower quartile were classified as Cold, and cells in the middle 50% were classified as Intermediate. These categories were used as local spatial ecosystem states and not as whole-tumor labels. A single tissue section could therefore contain Hot, Intermediate, and Cold tumor-enriched epithelial neighborhoods simultaneously. This classification generated 139,717 Hot cells, 139,717 Cold cells, and 279,433 Intermediate cells across the 17 HNSCC sections.

### 2.5 Pathway and module scoring

Pathway and module scores were quantified at single-cell resolution using curated transcript signatures represented within the Xenium panel(43). The analyzed modules included T-cell inflammatory signature score, IFN/antigen-presentation signature score, lipid metabolism score, prostaglandin/eicosanoid signaling score, cholesterol/sterol metabolism score, oxidative-redox stress score, fatty-acid/adipocyte-associated program score, NRF2-associated transcript score, ferroptosis-associated transcript score, mitochondrial stress/UPRmt signature score, radiation-stress signature score, EMT/invasion score, and NETosis signature score. For each module, the pathway score was calculated as the mean expression of available genes in the corresponding signature(20, 44). Because the Xenium panel is targeted, these scores were interpreted as transcript-derived program scores. They were not interpreted as direct measurements of metabolic flux, lipid mediator production, reactive oxygen species, NRF2 protein activation, mitochondrial function, ferroptotic cell death, or radiation-stress response.

### 2.6 Hot-minus-Cold pathway comparisons

To identify transcript programs associated with immune-hot tumor-enriched epithelial regions, mean pathway scores were compared between Hot and Cold tumor-enriched epithelial cells. For each pathway, the Hot-minus-Cold difference was calculated as:

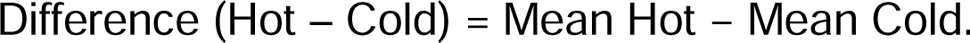

Positive values indicated higher transcript-derived pathway scores in Hot cells, whereas negative values indicated higher scores in Cold cells. Comparisons were performed across the full tumor-enriched epithelial cohort and also within each HNSCC section to evaluate whether pathway enrichment was reproducible across specimens. Two-sided Mann-Whitney U tests were used for group comparisons. Also, because cells are nested within tissue sections and sections are nested within tumors, cell-level statistical tests were interpreted descriptively. Biological conclusions were based on effect sizes, directionality, and consistency across HNSCC sections rather than nominal cell-level P values alone.

### 2.7 Metabolic ecosystem-state analysis

To examine whether tumor-enriched epithelial cells formed broader metabolic and stress-response states beyond the Hot/Cold classification, standardized pathway module scores were analyzed jointly across tumor-enriched epithelial cells. This analysis grouped tumor-enriched epithelial cells into six metabolic ecosystem states, following single-cell tumor ecosystem and metabolic-program scoring concepts that support transcriptomic identification of cell-state and metabolic heterogeneity within the tumor microenvironment(45). State-level module z-scores were then calculated to identify relative enrichment of inflammatory, lipid, eicosanoid, cholesterol, redox, ferroptosis-associated transcript scores, mitochondrial-stress, radiation-stress, EMT, and NETosis programs(44). The distribution of metabolic ecosystem states was compared across Hot, Cold, and Intermediate cells(32, 35). This analysis was used to determine whether immune-hot tumor-enriched epithelial cells represented a single uniform state or a collection of metabolically and stress-adapted sub-states(45, 46).

### 2.8 Spatial neighborhood analysis

Spatial neighborhood analyses were performed using Xenium-derived cell coordinates. For each tumor-enriched epithelial cell, nearby cells were grouped into immune neighbors, tumor neighbors, other non-tumor neighbors, and all neighboring cells. Mean pathway module scores were calculated within each neighboring compartment. Neighborhood pathway scores surrounding Hot and Cold tumor-enriched epithelial regions were then compared to determine whether immune-hot tumor-enriched epithelial regions were embedded within local environments enriched for inflammatory, metabolic, redox, mitochondrial-stress, or ferroptosis-associated programs. These neighborhood results were interpreted as spatial associations and not as proof of direct signaling, physical cell-cell contact, or causal cell-cell communication(46, 47).

### 2.9 Immune-Metabolic-Redox Ecosystem Score

An Immune-Metabolic-Redox Ecosystem Score, abbreviated IMRES, was developed to integrate the biological features that emerged from the Hot/Cold, metabolic, and neighborhood analyses. IMRES was designed around eleven candidate transcript-derived components: T-cell inflammatory signature score, CD274 expression, IFN/antigen-presentation score, lipid metabolism, prostaglandin/eicosanoid signaling, cholesterol metabolism, oxidative-redox lipid stress, NRF2-associated transcript score, ferroptosis-associated transcript score, mitochondrial stress, and radiation-stress response. Candidate component genes, Xenium-available genes, and missing genes are listed in **Supplementary Table S1**. The final primary IMRES score was calculated only from component scores with non-zero variance across tumor-enriched epithelial cells(41, 45, 48–51). Each retained component was z-score standardized across tumor-enriched epithelial cells. IMRES was calculated as the mean of the retained standardized components. Higher IMRES values reflected coordinated enrichment of immune activation, CD274-associated checkpoint biology, lipid and cholesterol remodeling, oxidative-redox adaptation, ferroptosis-associated programs, and radiation-stress responses(41, 44).

### 2.10 Tumor-immune proximity analysis

Tumor-immune proximity was quantified by nearest-neighbor analysis using Xenium spatial coordinates. For each tumor-enriched epithelial cell, the Euclidean distance to the nearest immune-associated cell was calculated(52). These distance measures contributed to Composite Hotness scoring and were also used to study spatial gradients, ecosystem-state organization, and later validation of Hot+IMRES_high_ tumor-enriched epithelial states. Distance values reflect coordinate-derived spatial proximity within the Xenium tissue plane(7, 46, 47). They should not be interpreted as evidence of immune synapse formation, tumor-cell killing, or direct membrane contact.

### 2.11 IMRES-associated gene analysis

To characterize the transcriptional features associated with IMRES, Spearman rank correlations were calculated between IMRES values and expression of all genes represented within the Xenium panel(44, 53). Genes were ranked according to correlation strength. Positively associated genes were used to identify markers of the immune-metabolic-redox ecosystem, while negatively associated genes were used to describe lower-IMRES (IMRES_low_) tumor-enriched epithelial states. This analysis was used for biological interpretation and signature development. It was not used to infer gene-regulatory causality(53).

### 2.12 Definition of Hot+IMRES_high_ ecosystem states

To resolve heterogeneity within Hot and Cold tumor-enriched epithelial cells, Composite Hotness classification was integrated with IMRES, following single-cell ecosystem-state and pathway-scoring frameworks used to define biologically distinct tumor and metabolic programs(35, 41, 44, 45). Tumor-enriched epithelial cells were first classified as Cold, Intermediate, or Hot based on Composite Hotness. The cohort-wide median IMRES value, calculated across all 558,867 tumor-enriched epithelial cells, was then used to define IMRES_high_ and IMRES_low_ populations. Hot and Cold tumor cells were subsequently subdivided according to IMRES status, generating four ecosystem states: Cold+IMRES_low_, Cold+IMRES_high_, Hot+IMRES_low_, and Hot+IMRES_high_. Intermediate cells were retained as a separate category and were not subdivided by IMRES. This approach resulted in five final ecosystem states: Cold+IMRES_low_, Cold+IMRES_high_, Intermediate, Hot+IMRES_low_, and Hot+IMRES_high_. The Hot+IMRES_high_ state was defined as tumor-enriched epithelial cells that were classified as both Hot by Composite Hotness and IMRES_high_ by the cohort-wide median threshold. Because this state exhibited the strongest immune, checkpoint-associated, metabolic, redox, and stress-response activity, it was used to represent the primary immune-metabolic-redox ecosystem in subsequent analyses.

### 2.13 Spatial permutation testing

A permutation-based spatial null model was used to test whether Hot+IMRES_high_ cells occupied non-random spatial positions relative to immune cells, consistent with spatial-omics and tumor-microenvironment approaches that compare observed cell-state proximity with randomized label assignments or null spatial distributions(46, 52, 54). Within each HNSCC section, Hot+IMRES_high_ labels were randomly reassigned while preserving the observed number of Hot+IMRES_high_ cells. Mean tumor-immune distances were recalculated across 500 permutations to generate null distributions. Observed tumor-immune distances for Hot+IMRES_high_ cells were then compared with the corresponding null distributions. This analysis tested whether Hot+IMRES_high_ cells were closer to immune-associated cells than expected by random label assignment within each tissue section.

### 2.14 Rank-based IMRES robustness analysis

To test whether the IMRES phenotype was robust to an alternative scoring strategy, a rank-based method was implemented. This approach was conceptually similar to AUCell-style single-cell gene-set scoring(43, 53), where gene-set signature score is estimated from within-cell gene rankings rather than absolute expression levels(43). AUCell and related rank-based methods are commonly used to score gene signatures in single-cell data because they evaluate each cell individually and are less dependent on normalization scale than expression-magnitude-based approaches(43, 44). Genes were first ranked according to their Spearman correlation with primary IMRES, following correlation-based single-cell feature-selection logic(53). The top 50 positively correlated genes and top 50 negatively correlated genes were used to construct positive and negative IMRES signatures. The complete Rank-IMRES_Positive_ and Rank-IMRESNegative gene lists are provided in **Supplementary Table S2**. For each tumor-enriched epithelial cell, within-cell gene ranks were used to calculate positive and negative rank-based signature scores. A net rank-based IMRES score was then calculated as:

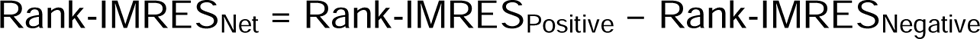

The relationship between rank-based IMRES and primary IMRES was assessed using Spearman correlation.

### 2.15 Benchmarking of IMRES and component scores

To compare how integrated and individual component scores related to the Hot+IMRES_high_ ecosystem, several candidate metrics were benchmarked. These included Composite Hotness Score, IMRES, rank-based IMRES, T-cell inflammatory signature score, CD274 expression, IFN/antigen-presentation score, lipid metabolism score, NRF2-associated transcript score, and ferroptosis-associated transcript score. Performance was assessed by receiver operating characteristic analysis, area under the receiver operating characteristic curve, area under the precision-recall curve, Mann-Whitney U testing, score-gene correlation benchmarking, and principal component analysis of IMRES-associated genes (36, 44, 55). These analyses were used to compare how integrated and single-component scores related to the predefined Hot+IMRES_high_ ecosystem state.

### 2.16 Statistical analysis and visualization

Statistical analyses were performed in Python using SciPy and scikit-learn(55). Comparisons between groups were evaluated using two-sided Mann– Whitney U tests, and associations between continuous variables were assessed using Spearman rank correlation coefficients. Classification performance was evaluated using receiver operating characteristic (ROC) analysis, area under the receiver operating characteristic curve (AUROC), and area under the precision-recall curve (AUPRC). Principal component analysis (PCA) was used for dimensionality reduction and visualization of IMRES-associated gene signatures. For spatial permutation analyses, observed tumor–immune distance measurements were compared with sample-specific null distributions generated by random reassignment of ecosystem labels while preserving the number of cells within each state. Correlation-based feature selection was performed by ranking genes according to Spearman correlation with IMRES, and rank-based robustness analyses were conducted using the top positively and negatively correlated genes. Data visualization was performed using Matplotlib and Seaborn and included heatmaps, boxplots, violin plots, bar plots, scatter plots, correlation matrices, network diagrams, PCA plots, ROC curves, ecosystem composition plots, and spatial summary visualizations. Unless otherwise specified, graphical summaries display median values and interquartile ranges. Because of the large number of cells analyzed, statistical significance was interpreted together with effect size, consistency across specimens and sections, and reproducibility across complementary analytical approaches. Statistical significance was defined as P < 0.05.

## 3. Results

### 3.1 Spatial classification identifies heterogeneous immune-hot and immune-cold tumor-enriched epithelial regions across HNSCC specimens

To characterize spatial immune heterogeneity across the GSE300147 cohort, 1,148,244 Xenium-profiled cells from 17 HNSCC tissue sections were analyzed after exclusion of the non-HNSCC ameloblastoma specimen. Tumor-enriched epithelial cells (n = 558,867) were classified using a Composite Hotness framework integrating T-cell inflammatory signature score, checkpoint signaling, CD274 expression, IFN/antigen-presentation programs, and tumor–immune proximity (**Fig. 1A**). Cell-level Composite Hotness scores showed clear separation of Cold, Intermediate, and Hot states (**Fig. 1B**). Across the cohort, 139,717 tumor-enriched epithelial cells were classified as Hot, 139,717 as Cold, and 279,433 as Intermediate. Sample-level composition varied substantially among specimens (**Fig. 1D**). The proportion of Hot tumor-enriched epithelial cells ranged from 1.17% in GSM9054485 to 52.08% in GSM9054477, whereas Cold-cell frequencies ranged from 6.03% to 78.79% across sections. The individual components contributing to the Composite Hotness score displayed coordinated differences across immune states (**Fig. 1C**). Hot tumor-enriched epithelial cells exhibited higher T-cell inflammatory, checkpoint, CD274, and IFN/antigen-presentation signature scores, whereas tumor–immune distance showed the opposite trend. Correlation analysis demonstrated that Composite Hotness was most strongly associated with T-cell inflammatory signature score (ρ = 0.781), IFN/AP signaling (ρ = 0.679), and checkpoint signaling (ρ = 0.670), while showing a strong inverse association with tumor–immune distance (ρ = −0.613) (**Fig. 1E**). Collectively, these findings support a continuum linking immune activation and tumor–immune proximity within HNSCC tissues.

**Figure 1.**
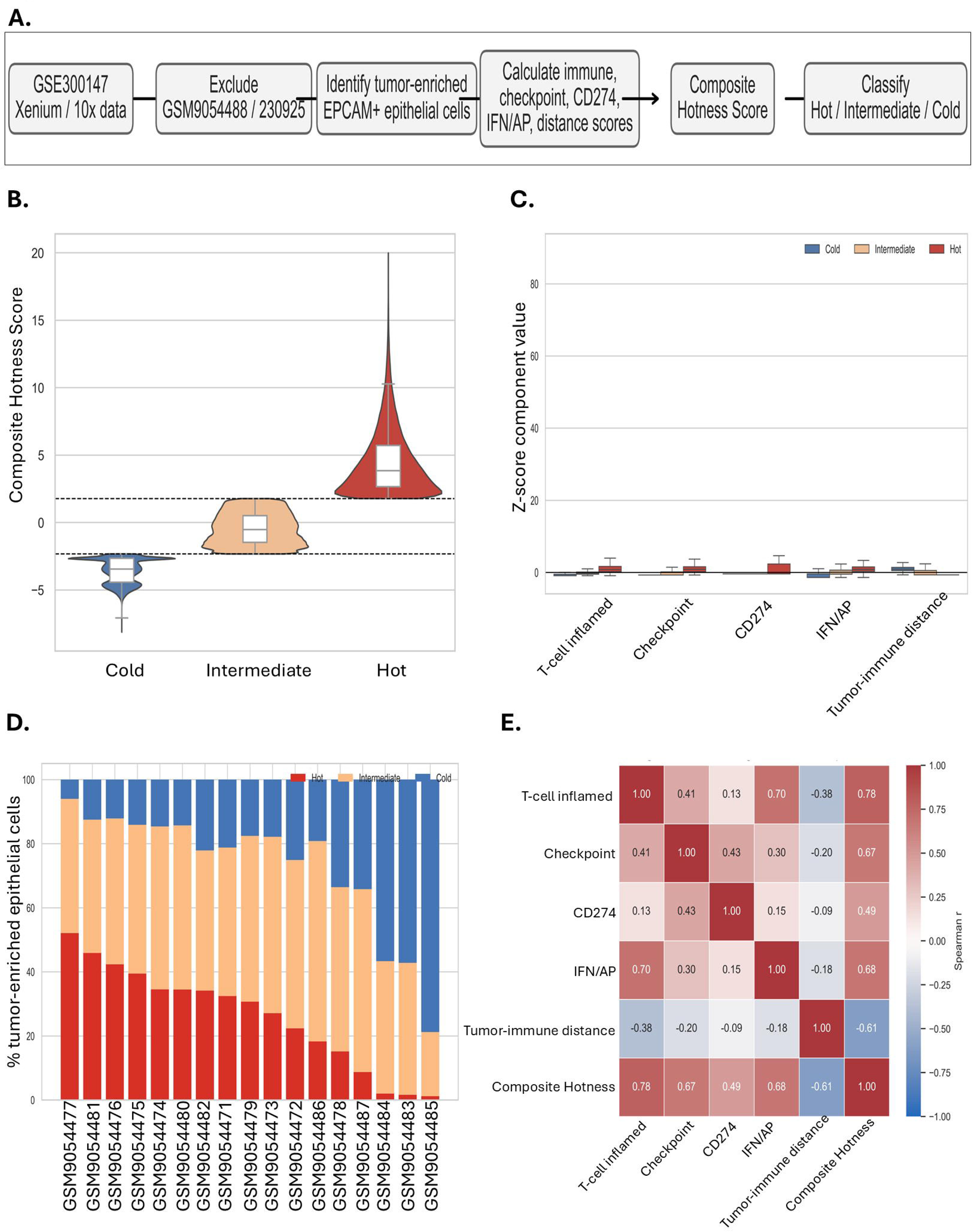
Identification of immune-hot and immune-cold tumor-enriched epithelial states in HNSCC. **(A)** Composite Hotness scoring framework integrating T-cell inflammatory signature score, checkpoint signaling, CD274 expression, IFN/antigen-presentation signature score, and tumor– immune proximity. **(B)** Composite Hotness score distribution across Cold, Intermediate, and Hot tumor-enriched epithelial states. **(C)** State-level z-score values of individual Composite Hotness components across Cold, Intermediate, and Hot tumor-enriched epithelial states. **(D)** Percentage composition of Hot, Intermediate, and Cold tumor-enriched epithelial cells across the 17 HNSCC sections. **(E)** Spearman correlation matrix of Composite Hotness components. Tumor-enriched epithelial cells were defined as EpCAM^+^ cells from the 17 HNSCC Xenium sections after exclusion of the non-HNSCC ameloblastoma specimen. Higher Composite Hotness scores indicate greater immune engagement and tumor–immune proximity.

### 3.2 Immune-hot tumor-enriched epithelial compartments occupy a distinct transcriptional state characterized by inflammatory and checkpoint-associated programs

Dimensionality-reduction analyses were performed to determine whether Hot and Cold tumor-enriched epithelial states occupied distinct transcriptional spaces. Principal component analysis and UMAP visualization revealed clear separation of Hot and Cold populations, with Intermediate cells occupying transitional positions between these extremes (**Fig. 2A-B**). Projection of Composite Hotness scores onto the UMAP embedding demonstrated a continuous gradient from immune-cold to immune-hot states across the transcriptional landscape (**Fig. 2C**). Sample-level centroid analyses further showed that Hot, Intermediate, and Cold populations were distributed across multiple HNSCC specimens, supporting that the observed transcriptional organization was not dominated by a single section (**Fig. 2F**). Marker-gene heatmaps demonstrated enrichment of immune activation, cytotoxicity, checkpoint, and antigen-presentation programs within the Hot compartment, including increased expression of PTPRC, CD3D, CD3E, CD8A, PRF1, GZMB, CXCL9, CXCL10, CCL5, PDCD1, CTLA4, TIGIT, and HLA-DRA (**Fig. 2D**). Because EpCAM^+^ was used as an enrichment strategy and Xenium segmentation assigns transcripts within spatial compartments, immune-associated transcripts in this compartment were interpreted as tumor-neighborhood or tumor-associated signals rather than definitive tumor-cell-intrinsic immune gene expression. Component-level analysis further confirmed progressive enrichment of immune-related features across the Cold-to-Hot continuum (**Fig. 2E**). T-cell inflammatory signature score increased from −1.04 in Cold cells to 1.35 in Hot cells; checkpoint signaling increased from −0.93 to 1.39; CD274 increased from −0.84 to 1.41; IFN/AP increased from −1.25 to 1.20; tumor-immune distance declined from 1.38 to −0.97 (**Fig. 2E**). Together, these analyses indicate that immune-hot tumor-enriched epithelial regions represent a distinct tumor-enriched epithelial/transcriptomic neighborhood state rather than only reflecting increased immune-cell abundance.

**Figure 2.**
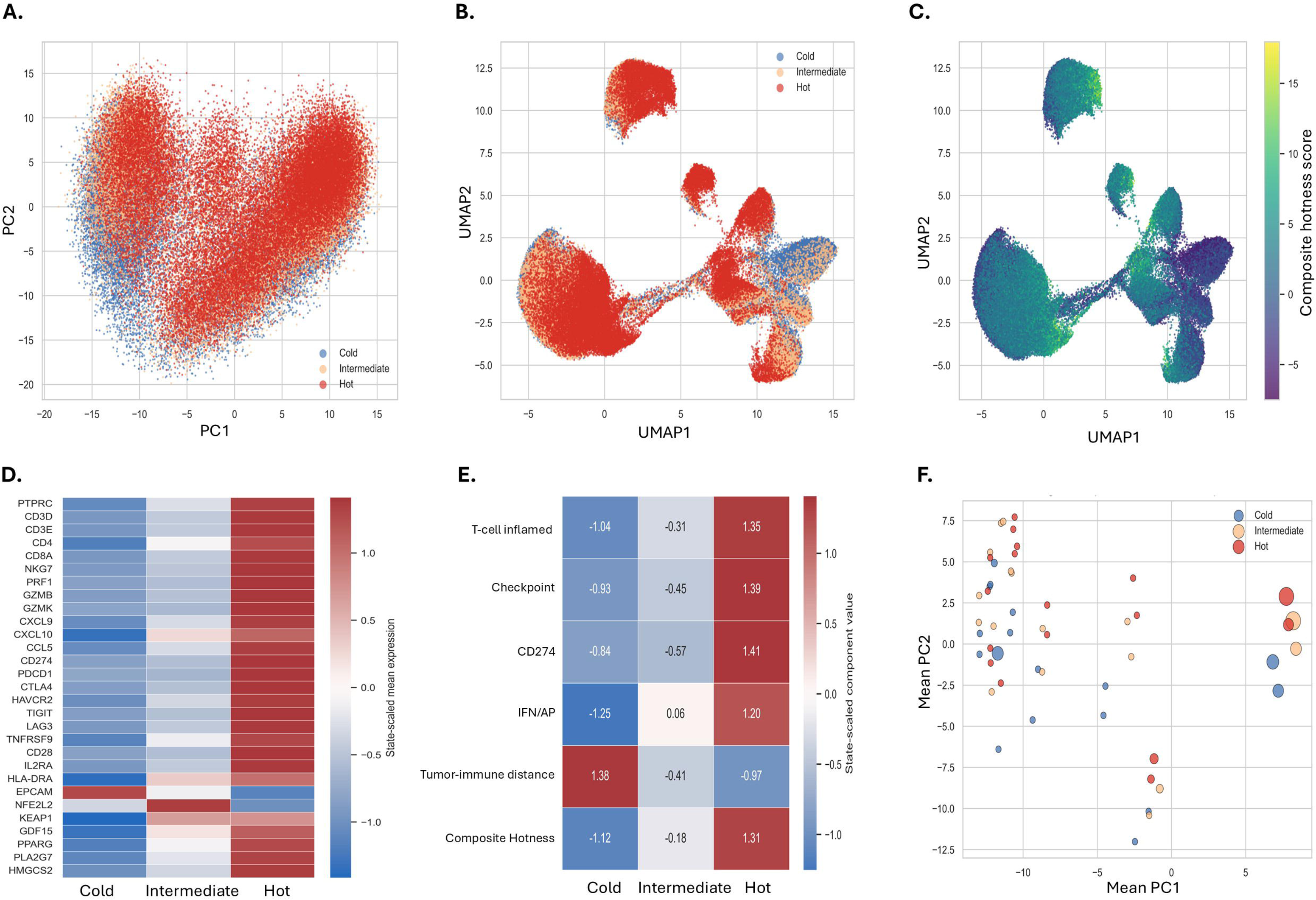
Transcriptomic organization of immune ecosystem states in HNSCC. **(A)** Principal component analysis (PCA) of tumor-enriched epithelial cells colored by immune ecosystem state. **(B)** UMAP visualization of Cold, Intermediate, and Hot tumor-enriched epithelial populations. **(C)** Projection of Composite Hotness scores onto the UMAP embedding. **(D)** Heatmap of representative immune, cytotoxic, checkpoint, and antigen-presentation markers across ecosystem states. **(E)** Standardized Composite Hotness component scores across Cold, Intermediate, and Hot populations. **(F)** Sample-level state centroids plotted by mean PC1 and mean PC2 values, demonstrating that Cold, Intermediate, and Hot ecosystem states are represented across multiple HNSCC specimens. Heatmaps display state-level standardized expression or transcript-derived pathway scores.

### 3.3 Immune-hot tumor-enriched epithelial regions are associated with coordinated metabolic and stress-response ecosystems

To determine whether immune activation was accompanied by broader biological changes, transcript-derived metabolic and stress-response programs were quantified in tumor-enriched epithelial cells. Multiple pathways were enriched in Hot tumor-enriched epithelial cells relative to Cold cells, including lipid metabolism (Δ = 0.033), cholesterol/sterol metabolism (Δ = 0.082), and fatty-acid/adipocyte-associated programs (Δ = 0.100) (**Fig. 3A-B**). More substantial differences were observed in oxidative-redox stress score (Δ = 0.288), NRF2-associated transcript score (Δ = 0.252), ferroptosis-associated transcript score (Δ = 0.215), mitochondrial stress/UPRmt signature score (Δ = 0.431), and radiation-stress signature score (Δ = 0.332), with all comparisons remaining highly significant (**Fig. 3B**). Among all modules evaluated, T-cell inflammatory signature score exhibited the largest Hot-minus-Cold difference (Δ = 1.15) (**Fig. 3B**). These patterns were broadly conserved across the 17 HNSCC sections, indicating that the observed metabolic and stress-associated features were not driven by a small subset of samples (**Fig. 3C**). Extension of these biological programs into the surrounding microenvironment was then evaluated. Compared with Cold regions, Hot tumor-enriched epithelial regions were associated with neighboring cells displaying higher T-cell inflammatory signature score (Δ = 0.621), oxidative-redox stress score (Δ = 0.140), NRF2-associated transcript score (Δ = 0.111), ferroptosis-associated transcript score (Δ = 0.084), and radiation-stress signature score (Δ = 0.210) (**Fig. 3D**). Similar trends were observed across immune-cell, tumor-enriched epithelial, and non-tumor neighboring compartments. Immune-cell neighborhoods surrounding Hot tumor-enriched epithelial cells showed particularly strong enrichment for T-cell inflammatory signature score (Δ = 0.663), and radiation-stress programs (Δ = 0.165), while non-tumor neighboring compartments displayed marked enrichment of oxidative-redox stress, NRF2-associated transcript score, ferroptosis-associated programs, and radiation-stress responses (**Fig. 3D**). Together, these findings indicate that immune-hot tumor-enriched epithelial regions are embedded within broader metabolic and stress-associated ecosystems rather than representing isolated tumor-cell states.

**Figure 3.**
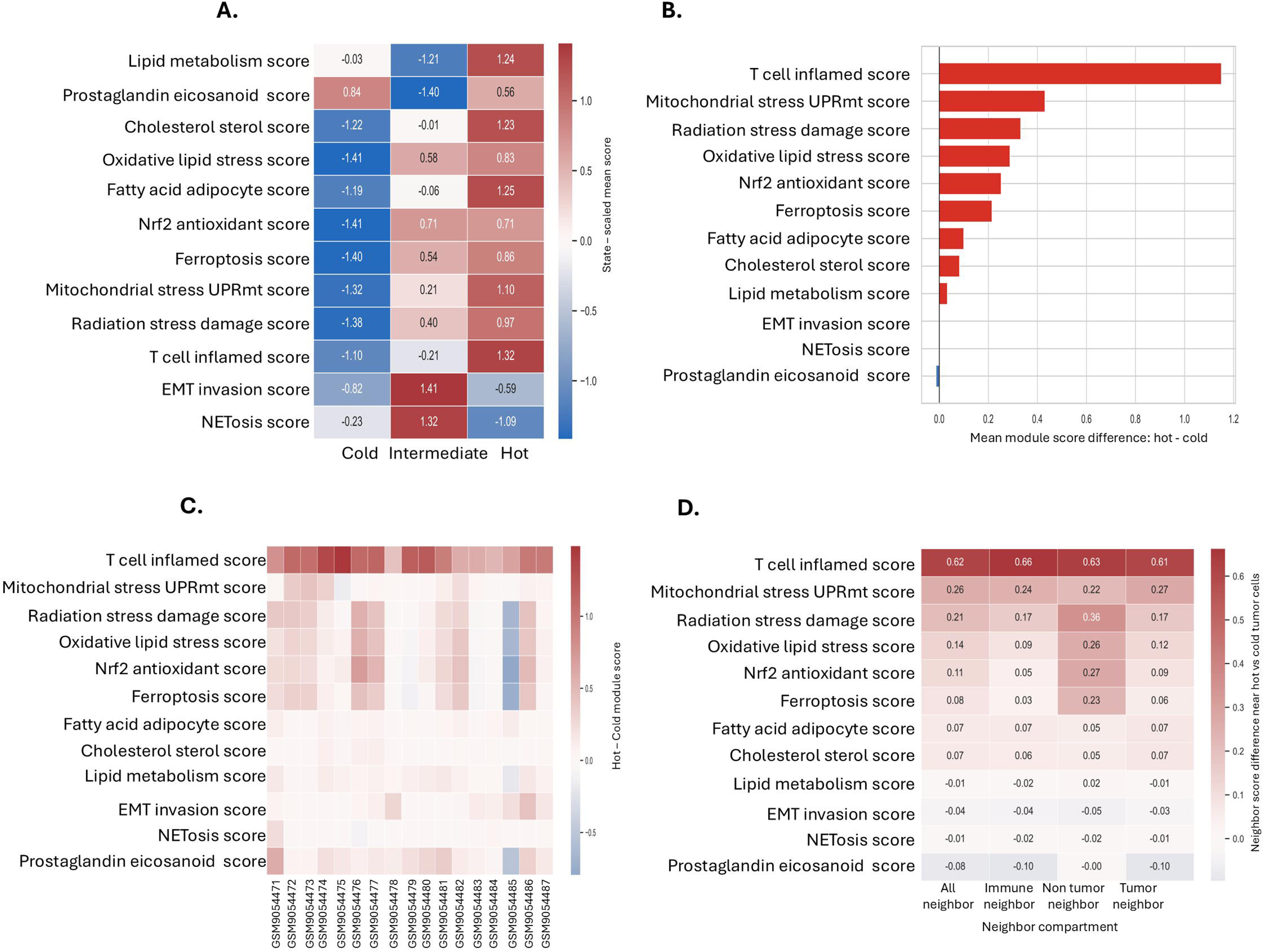
Metabolic and stress-response programs associated with immune ecosystem states. **(A)** Mean transcript-derived pathway scores across Cold, Intermediate, and Hot tumor-enriched epithelial states. **(B)** Hot-minus-Cold differences for immune, metabolic, redox, ferroptosis-associated, mitochondrial-stress, and radiation-stress programs. **(C)** Sample-level conservation of Hot-minus-Cold pathway differences across HNSCC sections. **(D)** Neighborhood pathway analysis comparing cells surrounding Hot and Cold tumor-enriched epithelial regions. Pathway scores were calculated from transcript-derived signatures. Neighborhood analyses were performed using local cellular compartments surrounding tumor-enriched epithelial cells. All pathway scores are transcript-derived signature scores and do not represent direct measurements of protein abundance, metabolic flux, oxidative stress, ferroptosis, or mitochondrial function.

### 3.4 Metabolic ecosystem states reveal coordinated immune, metabolic, and stress-response niches within HNSCC tumors

To determine whether the metabolic and stress-response programs associated with immune-hot and immune-cold tumor-enriched epithelial cells reflected broader biological organization, tumor-enriched epithelial cells were grouped according to transcript-derived metabolic signatures. This analysis identified six recurrent ecosystem states with distinct functional profiles (**Fig. 4A-B**). Lipid-, cholesterol-, and fatty-acid–associated programs were concentrated within specific ecosystem states, whereas oxidative-redox stress, ferroptosis-associated transcript score, mitochondrial stress, and radiation-stress responses clustered within separate states, indicating that these biological processes are not uniformly distributed across tumors (**Fig. 4A**). Importantly, multiple ecosystem states were present within both Hot and Cold tumor-enriched epithelial populations (**Fig. 4B**), demonstrating that immune activation alone does not fully explain the metabolic diversity observed across tumor-enriched epithelial cells. Although tumor-immune distance was evaluated as a potential spatial feature (**Fig. 4C**), uneven representation across distance bins limited interpretation of distance-dependent pathway gradients. To assess reproducibility, pathway enrichment patterns were examined across all 17 HNSCC specimens (**Fig. 4D**). T-cell inflammatory signature score was consistently enriched in Hot regions in every tumor analyzed (17/17 samples; mean Hot-minus-Cold difference = 0.95). Fatty-acid/adipocyte-associated programs were similarly enriched across all specimens, while lipid metabolism, prostaglandin/eicosanoid signaling, EMT-associated programs, mitochondrial stress, oxidative-redox stress, NRF2-associated transcript score, ferroptosis-associated transcript score, and radiation-stress responses remained positively enriched in most tumors (**Fig. 4D**). These findings indicate that the observed ecosystem structure is broadly conserved across the cohort rather than driven by a small subset of samples.

**Figure 4.**
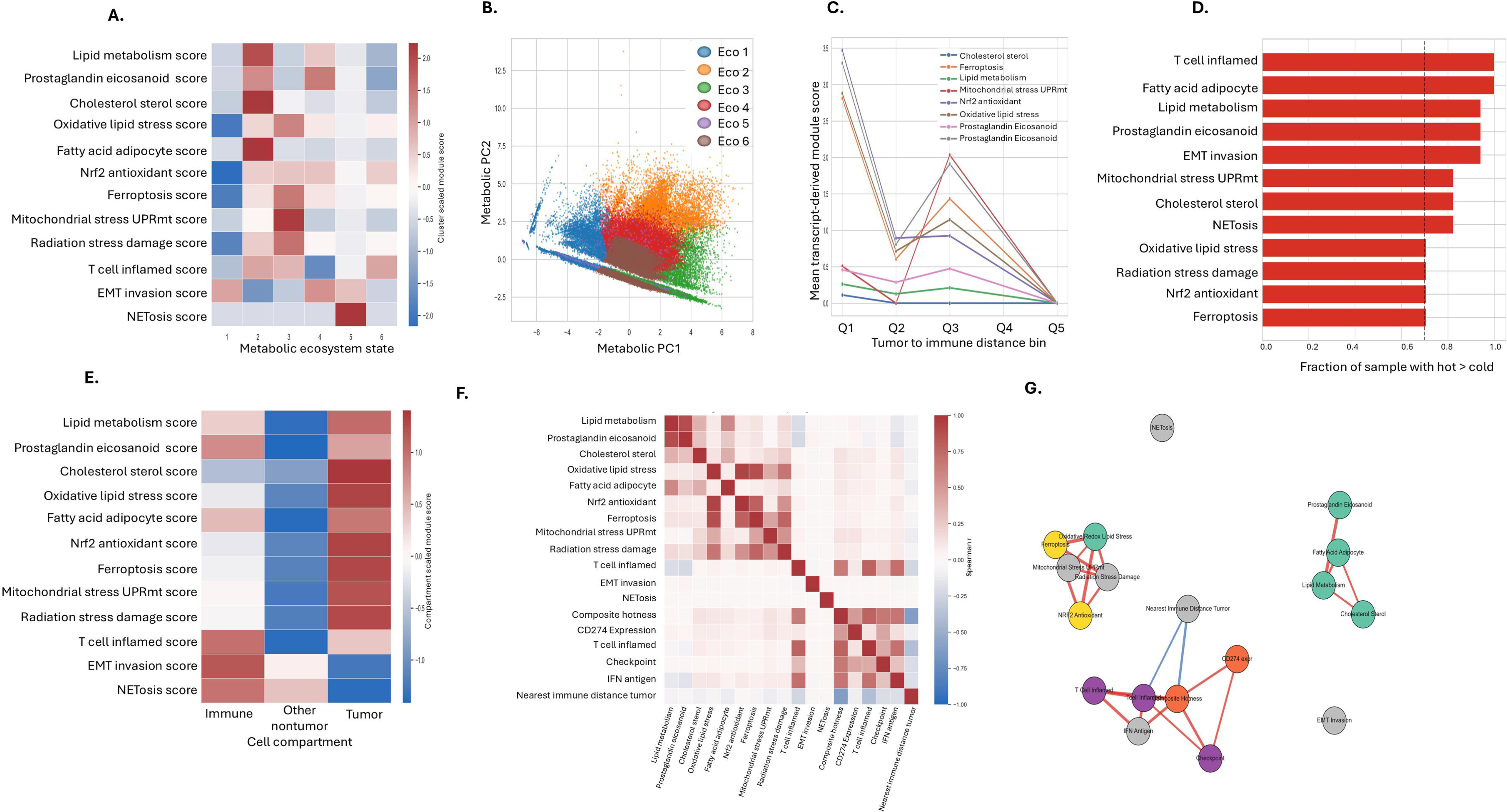
Spatial immune-metabolic-redox ecosystem analysis. **(A)** Pathway enrichment across six metabolic ecosystem states, with putative functional profiles inferred from dominant transcript-derived module scores: Eco1, EMT-high/metabolic-redox-low; Eco2, lipid-eicosanoid/cholesterol-high; Eco3, redox-ferroptosis/stress-high; Eco4, eicosanoid-EMT/immune-low; Eco5, NETosis-high; and Eco6, T-cell-inflamed/lipid-low. **(B)** Two-dimensional embedding of six metabolic ecosystem states derived from pathway-score profiles. **(C)** Exploratory pathway summaries across tumor-immune distance bins. **(D)** Conservation of pathway enrichment patterns across the 17 HNSCC sections. **(E)** Transcript-derived pathway scores within immune, tumor-enriched epithelial, and non-tumor cellular compartments. **(F)** Correlation matrix of pathway scores. **(G)** Network representation of significant pathway associations. Network edges represent significant positive pathway correlations. Tumor-immune distance analyses are presented as exploratory.

Analysis of cellular compartments revealed distinct biological specializations (**Fig. 4E**). Tumor-enriched epithelial cells showed the strongest enrichment for cholesterol metabolism, oxidative-redox stress, NRF2-associated transcript score, ferroptosis-associated programs, mitochondrial stress, and radiation-stress signature score, whereas immune-cell compartments were preferentially associated with T-cell inflammation, EMT-associated programs, and NETosis signatures. Correlation and network analyses showed that oxidative-redox stress, NRF2-associated transcript score, ferroptosis-associated transcript score, mitochondrial stress, and radiation-stress programs formed a tightly connected module (**Fig. 4F-G**). Notably, oxidative-redox stress score was strongly correlated with NRF2-associated transcript score (ρ = 0.936) and ferroptosis-associated transcript score (ρ = 0.876), supporting the existence of an integrated stress-adaptation network within HNSCC tumors. Together, these analyses identify a reproducible immune-metabolic-redox ecosystem architecture that extends beyond simple Hot-versus-Cold immune classification and is conserved across HNSCC specimens.

### 3.5 Development of an Immune-Metabolic-Redox Ecosystem Score (IMRES)

The coordinated enrichment of immune, metabolic, redox, and stress-response pathways in immune-hot tumor-enriched epithelial regions prompted the development of an integrated Immune-Metabolic-Redox Ecosystem Score (IMRES). IMRES was developed from candidate immune, metabolic, redox, and stress-response transcript features, with the final score calculated from the retained non-zero-variance components represented in the analyzed Xenium data (**Fig. 5A,G**). The primary IMRES component gene sets are listed in **Supplementary Table S1**. Across 558,867 tumor-enriched epithelial cells, IMRES showed a broad distribution and increased progressively from Cold to Intermediate to Hot tumor-enriched epithelial states (**Fig. 5A-B**). Mean IMRES values were −0.279 in Cold cells, −0.021 in Intermediate cells, and 0.320 in Hot cells, with all pairwise comparisons remaining highly significant. These results indicate that IMRES captures biological features associated with increasing immune engagement within the tumor microenvironment (**Fig. 5B,G**). Further, to identify molecular programs linked to IMRES, genes most strongly associated with the score were examined (**Fig. 5C-F**). The highest positive correlations were observed for NFE2L2 (r = 0.505), GDF15 (r = 0.472), HLA-DRA (r = 0.345), CD274 (r = 0.315), KEAP1 (r = 0.293), and MDM2 (r = 0.279). These genes connect higher IMRES values with oxidative-stress adaptation, cellular stress responses, antigen presentation, and checkpoint-associated biology. Because IMRES includes immune-related components as well as metabolic, redox, and stress-response components, part of its association with Composite Hotness is expected by design. The moderate correlation between IMRES and Composite Hotness suggests overlap but not identity; future analyses could further decompose immune and non-immune IMRES components (**Fig. 5I**). Across tumor-enriched epithelial cells, IMRES showed a moderate correlation with Composite Hotness (r = 0.48), indicating that immune-hot regions generally exhibit higher IMRES values while retaining substantial metabolic and stress-response heterogeneity. At the specimen level, elevated IMRES values were observed in Hot regions across most HNSCC sections (**Fig. 5H**). For example, mean IMRES increased from 0.226 to 0.825 in GSM9054474, from 0.236 to 0.750 in GSM9054475, and from −0.211 to 0.382 in GSM9054482. Although the magnitude of enrichment varied among tumors, the overall pattern remained consistent across the cohort. Collectively, these findings establish IMRES as an integrated measure of immune activation, metabolic remodeling, redox adaptation, ferroptosis-associated biology, and cellular stress responses. The strong association of IMRES with NFE2L2, GDF15, HLA-DRA, and CD274, together with its reproducible enrichment in immune-hot tumor-enriched epithelial regions, supports its use as a framework for defining immune-metabolic-redox ecosystem states in HNSCC.

**Figure 5.**
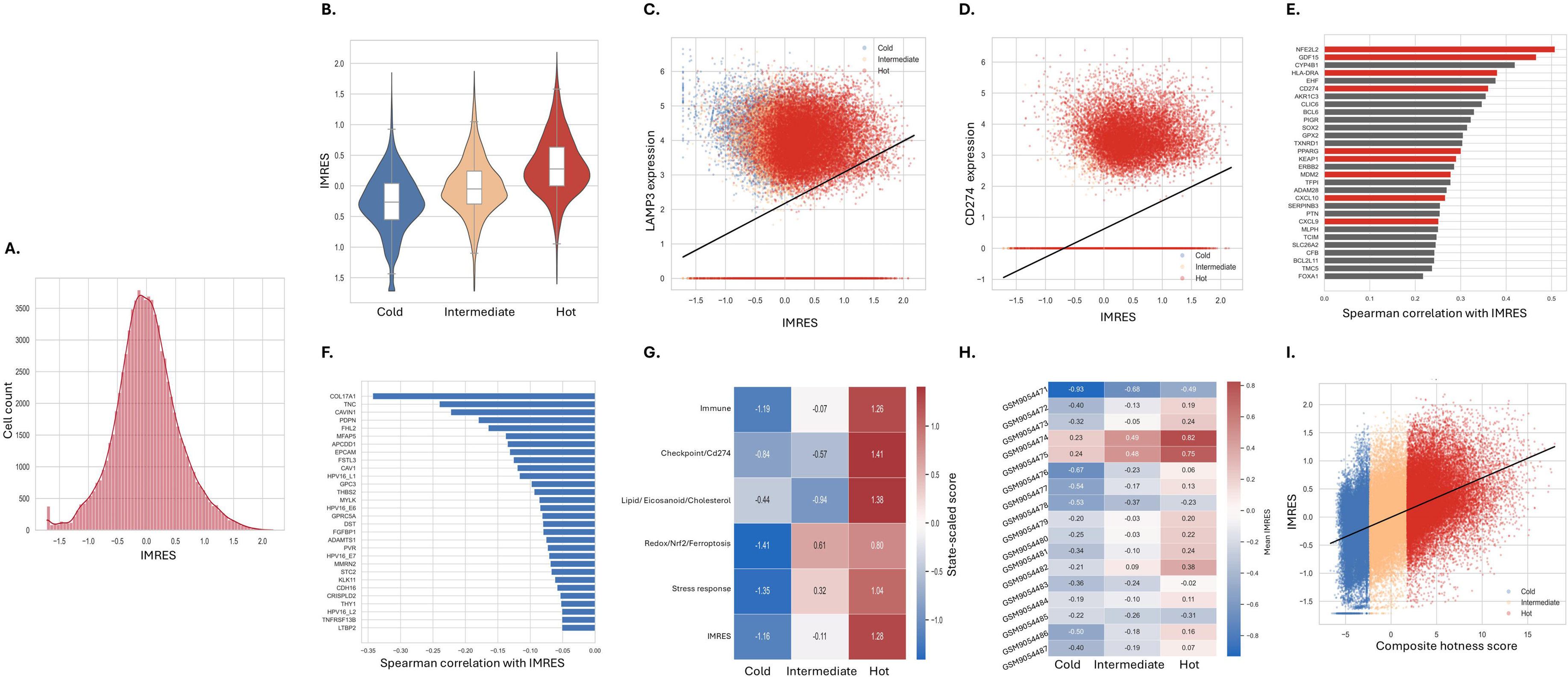
Development and characterization of the Immune-Metabolic-Redox Ecosystem Score (IMRES). **(A)** Distribution of IMRES across all tumor-enriched epithelial cells. **(B)** IMRES across Cold, Intermediate, and Hot immune ecosystem states. **(C-D)** Representative correlations between IMRES and two representative immune-metabolic-redox genes (LAMP3 and CD274 respectively). **(E)** Top positively IMRES-associated genes. **(F)** Top negatively IMRES-associated genes. **(G)** IMRES component structure across Cold, Intermediate, and Hot tumor states. **(H)** Sample-level IMRES values across HNSCC specimens. **(I)** Relationship between IMRES and Composite Hotness. “IMRES was designed from eleven candidate transcript-derived immune, metabolic, redox, and stress-response features, with the final score calculated from retained non-zero-variance components represented in the analyzed Xenium data. Positive IMRES values indicate stronger immune-metabolic-redox ecosystem activity. Correlations were calculated using all 558,867 tumor-enriched epithelial cells. Primary IMRES component gene sets are provided in **Supplementary Table S1**.

### 3.6 Hot+IMRES_high_ cells define a major inflamed and stress-adapted tumor-enriched epithelial ecosystem

Although IMRES increased across the Cold-to-Hot continuum, substantial heterogeneity remained within both Hot and Cold tumor-enriched epithelial populations. To resolve this variation, tumor-enriched epithelial cells were stratified using both Composite Hotness and IMRES, generating five Composite Hotness–IMRES ecosystem states: Cold+IMRES_low_ (99,143 cells), Cold+IMRES_high_ (40,574 cells), Intermediate (279,433 cells), Hot+IMRES_low_ (32,843 cells), and Hot+IMRES_high_ (106,874 cells) (**Fig. 6A**). The Hot+IMRES_high_ population represented the major immune-inflamed ecosystem subset among Hot cells, whereas most Cold tumor-enriched epithelial cells were classified as Cold+IMRES_low_. Comparison of molecular features across ecosystem states revealed a progressive increase in immune, metabolic, and stress-response scores, with the strongest enrichment observed in the Hot+IMRES_high_ compartment (**Fig. 6B**). Relative to Cold+IMRES_low_ cells, Hot+IMRES_high_ tumor-enriched epithelial cells showed higher T-cell inflammatory signature score (1.32 vs. 0.14), CD274 expression (1.88 vs. 0.00), IFN/antigen-presentation signaling (3.54 vs. 0.72), oxidative-redox stress score (3.23 vs. 2.33), NRF2-associated transcript score (3.81 vs. 2.90), ferroptosis-associated transcript score (3.13 vs. 2.30), and radiation-stress signature score (3.67 vs. 2.77). The largest differences were observed for redox adaptation, ferroptosis-associated programs, checkpoint-associated programs, and separately assessed mitochondrial-stress signatures, indicating that Hot+IMRES_high_ cells represent more than a highly inflamed state alone.

**Figure 6.**
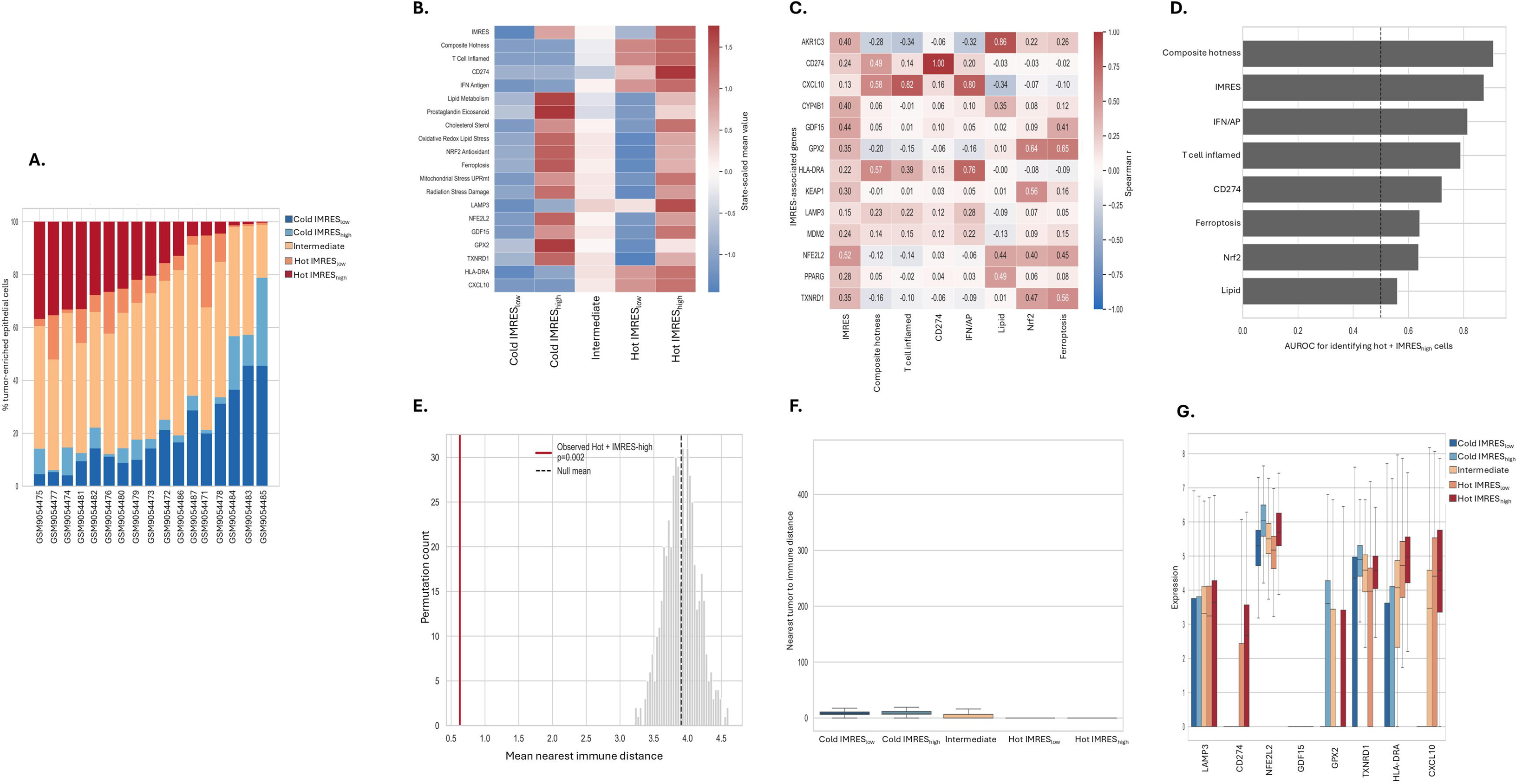
Biological and spatial characterization of the Hot+IMRES_high_ ecosystem state. **(A)** Sample-level distribution of Composite Hotness–IMRES ecosystem states. **(B)** Immune, metabolic, redox, and stress-response features across ecosystem states. **(C)** Correlation benchmark of candidate scores against representative ecosystem-associated genes. **(D)** AUROC benchmarking of candidate metrics for identification of Hot+IMRES_high_ cells. **(E)** Spatial permutation analysis comparing observed and randomized Hot+IMRES_high_ immune proximity. **(F)** Tumor-immune proximity across ecosystem states. **(G)** Expression of representative immune-, antigen-presentation-, and stress-adaptation-associated genes across ecosystem states. Ecosystem-state colors are maintained throughout the manuscript: Cold+IMRES_low_ (dark blue), Cold+IMRES_high_ (light blue), Intermediate (orange), Hot+IMRES_low_ (light red), and Hot+IMRES_high_ (dark red). Permutation testing was performed using randomized ecosystem labels while preserving ecosystem-state frequencies within each tissue section.

To determine whether IMRES captured information beyond individual pathways, its performance was compared with several candidate metrics (**Fig. 6C-D**). IMRES remained strongly associated with genes linked to immune activation and stress adaptation, including NFE2L2, GDF15, HLA-DRA, CD274, GPX2, TXNRD1, and CXCL10 (**Fig. 6C**). Benchmark analysis further showed that IMRES showed stronger discrimination than individual pathway scores such as NRF2-associated transcript score, ferroptosis-associated transcript score, lipid metabolism, and CD274 expression in identifying Hot+IMRES_high_ cells (AUROC = 0.874), although Composite Hotness showed the strongest discrimination because it contributed to the definition of the Hot+IMRES_high_ state (AUROC = 0.908) (**Fig. 6D**). These findings support the value of IMRES as an integrated measure of the immune-metabolic-redox ecosystem scores rather than a surrogate for any single biological feature. The Hot+IMRES_high_ state also showed distinct spatial organization. Permutation-based spatial modeling showed that Hot+IMRES_high_ cells occurred significantly closer to immune cells than expected by chance (observed mean distance = 0.629; null mean = 3.906; empirical P = 0.002) (**Fig. 6E**). Direct comparison across ecosystem states similarly demonstrated that Hot+IMRES_high_ cells occupied the most immune-proximal locations, whereas Cold+IMRES_low_ cells were more distant from immune populations (**Fig. 6F**). Consistent with this spatial pattern, the Hot+IMRES_high_ compartment displayed the highest expression of representative immune-metabolic-redox genes, including CD274, NFE2L2, GDF15, GPX2, TXNRD1, HLA-DRA, and CXCL10 (**Fig. 6G**). Together, these results identify Hot+IMRES_high_ as a distinct Composite Hotness–IMRES ecosystem state in which immune activation, cellular stress adaptation, and immune-cell proximity converge within the same tumor-enriched epithelial regions.

### 3.7 Rank-based scoring supports the robustness of the IMRES framework

To determine whether the IMRES phenotype depended on the original module-based scoring framework, an alternative rank-based score was generated using genes selected by correlation with the primary IMRES score (**Fig. 7A**). Among the positively associated genes were NFE2L2, GDF15, CYP4B1, HLA-DRA, CD274, AKR1C3, GPX2, TXNRD1, PPARG, KEAP1, MDM2, CXCL10, CXCL9, LAMP3, CD8A, CD3E, CD3D, CCL5, and PLA2G7, whereas negatively associated genes included COL17A1, TNC, CAVIN1, PDPN, FHL2, MFAP5, APCDD1, EpCAM, FSTL3, CAV1, FGFBP1, PVR, IL1RL1, and SNAI1. Using the top 50 positively and negatively associated genes, rank-based scores were calculated across 558,867 tumor-enriched epithelial cells (**Fig. 7B**). The full positive and negative gene sets used for rank-based scoring are provided in **Supplementary Table S2**. This alternative scoring strategy closely reproduced the original ecosystem structure, with Hot+IMRES_high_ cells consistently showing the highest rank-based scores and Cold+IMRES_low_ cells showing the lowest. Despite using a different scoring strategy, rank-based and module-based IMRES scores remained moderately correlated (Spearman r = 0.597; **Fig. 7C**). Additional benchmarking analyses showed that IMRES and rank-based IMRES performed comparably to established immune-related metrics in identifying Hot+IMRES_high_ cells (**Fig. 7D-E**). Both scoring approaches also maintained consistent associations with genes linked to immune activation, antigen presentation, checkpoint signaling, and oxidative-stress adaptation, including NFE2L2, GDF15, CD274, HLA-DRA, CXCL10, GPX2, TXNRD1, and KEAP1 (Fig. 7F). Principal component analysis provided an additional dimensionality-reduction view that recovered the same ecosystem organization, separating Hot+IMRES_high_ and Cold+IMRES_low_ populations while positioning intermediate states between these extremes (**Fig. 7G**). Together, **Figure 7** provides an internal robustness test of the manuscript’s central observation rather than independent external validation. Earlier analyses established that HNSCC tumors contain spatially organized tumor-enriched epithelial ecosystems associated with immune activation, antigen presentation, checkpoint signaling, metabolic adaptation, oxidative-stress responses, and cellular stress programs. Here, the same biological organization was recovered using correlation-based gene selection, an alternative rank-based scoring approach, benchmarking analyses, score-gene association testing, and dimensionality reduction. These results support Hot+IMRES_high_ as an analytically robust ecosystem state with coordinated immune, metabolic, and redox features. Because all analyses were derived from the same public Xenium resource, these findings should be interpreted as analytical robustness and internal reproducibility, not as independent cohort validation.

**Figure 7.**
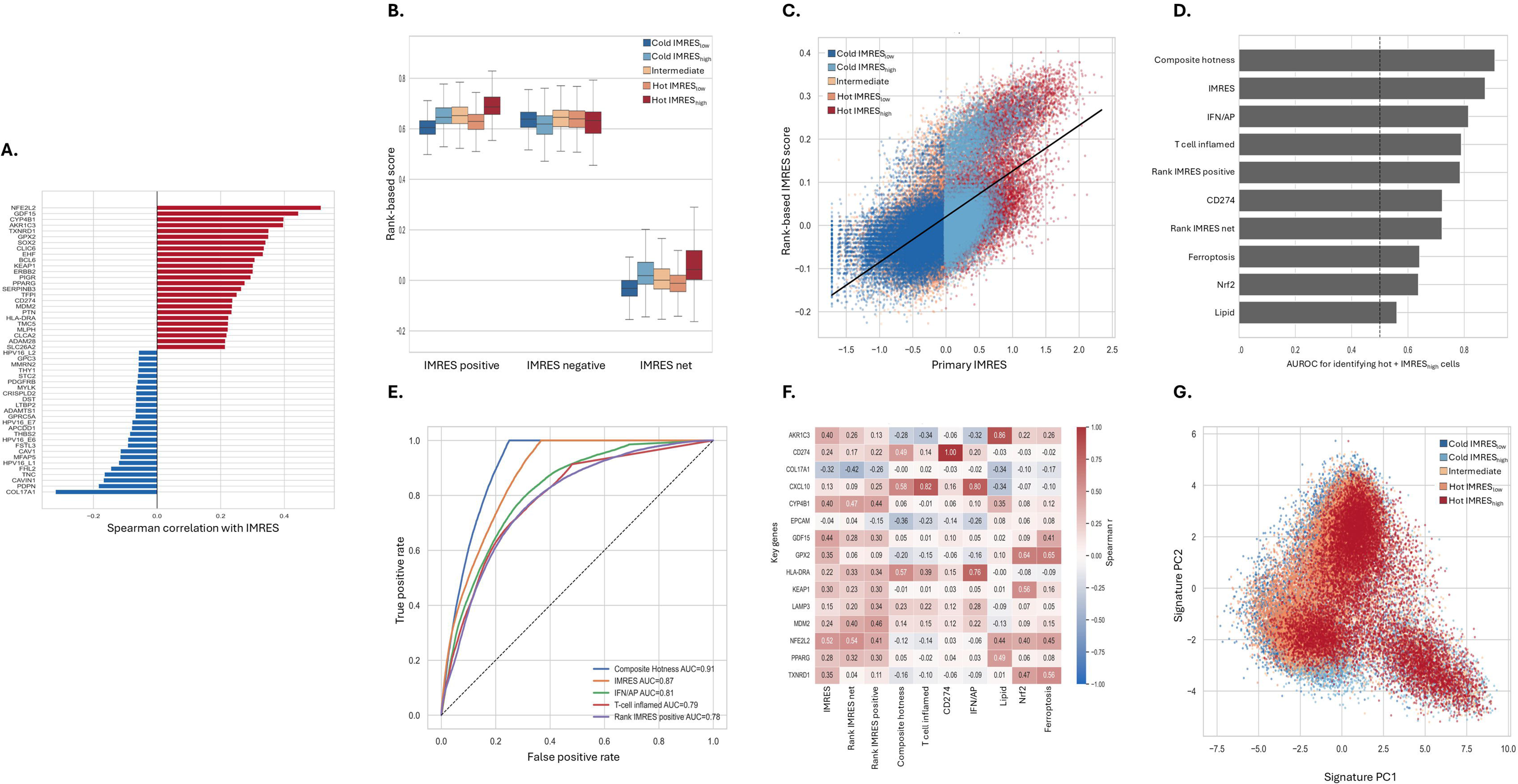
Rank-based robustness analysis of the IMRES ecosystem framework. **(A)** Correlation-based selection of IMRES-associated gene. **(B)** Rank-based IMRES component scores across ecosystem states. **(C)** Relationship between rank-based IMRES and module-based IMRES. **(D)** Benchmarking performance of candidate metrics for identifying Hot+IMRES_high_ cells. **(E)** Receiver operating characteristic curves. **(F)** Score-gene correlation benchmark across representative immune and stress-response genes. **(G)** Principal component analysis of top IMRES-associated genes. Rank-based IMRES was calculated using the top positively and negatively IMRES-correlated genes selected by correlation with the primary module-based IMRES score. Colors correspond to ecosystem-state definitions shown in Figure 6. Xenium genes/features used for Rank-IMRES_Positive_ and Rank-IMRES_Negative_ scoring are listed in **Supplementary Table S2**.

## 4. Discussion

The original GSE300147 study by McCord and colleagues(23) provided an important spatial T-cell framework for HNSCC by linking T-cell receptor clonality, transcriptional state, and spatial position within intact tumor tissues. Their work showed that presumed tumor-specific T cells are broadly distributed across the tumor microenvironment and that cells from the same clone could occupy different transcriptional states depending on local tissue context, with immune-rich and tumor-dense regions supporting different T-cell phenotypes. The present study builds on that resource but addresses a different question and inspected how tumor-enriched epithelial cells are organized across immune-hot and immune-cold spatial states and whether immune activation is coupled to metabolic, redox, and stress-adaptive programs(28). Thus, in this reanalysis, immune-hot HNSCC regions were not defined by immune-cell abundance alone, instead they marked spatially organized tumor-enriched epithelial neighborhoods where immune activation coincided with checkpoint-associated signaling, metabolic remodeling, redox adaptation, and stress-response programs. Hot, Intermediate, and Cold tumor-enriched epithelial states coexisted across the analyzed HNSCC sections, indicating that immune signature score is spatially heterogeneous within and across tumors. This is consistent with the idea that immune-cell abundance alone does not adequately describe the tumor immune microenvironment(6, 8, 12, 34).

Spatial transcriptomic studies and reviews have emphasized that cellular location, tissue architecture, cell-cell proximity, and local neighborhood structure can carry information that is lost in dissociated single-cell or bulk profiling approaches(8, 12, 34). In that context, a tumor-centered layer is added to the original T-cell-centered GSE300147 analysis by showing that epithelial tumor-enriched compartments also organize into spatially distinct immune states. The most important biological observation is that immune-hot tumor-enriched epithelial regions carried a coordinated stress-adaptive program(32, 35, 41, 45, 48). As expected, Hot cells showed higher T-cell inflammatory signature score, IFN/antigen-presentation signaling, checkpoint-associated score, and CD274 expression. However, the same regions also showed enrichment of lipid and cholesterol metabolism, oxidative-redox stress, NRF2-associated transcript score, ferroptosis-associated programs, mitochondrial stress, and radiation-stress signatures (41, 45, 49–51). This finding suggests that immune activation and stress adaptation may coexist within the same local tumor neighborhoods, rather than representing separate tumor states. This interpretation is consistent with recent HNSCC work showing that KEAP1/NFE2L2 pathway alterations are linked to treatment resistance and an unfavorable tumor-immune microenvironment, including radiotherapy resistance and suppressed antitumor immunity. It is also consistent with broader HNSCC literature connecting NRF2-associated programs to antioxidant defense, ferroptosis evasion, and therapeutic resistance(50, 56).

The development of IMRES was intended to capture this integrated biology in a single quantitative framework. IMRES increased across the Cold-to-Intermediate-to-Hot continuum and was most strongly associated with genes such as NFE2L2, GDF15, HLA-DRA, CD274, KEAP1, and MDM2, placing oxidative-stress adaptation, cellular stress signaling, antigen presentation, and checkpoint-associated biology within the same tumor-enriched epithelial ecosystem(41, 50, 56). This distinction is important because an immune-hot region may not necessarily be immune-permissive. Instead, immune activation may occur alongside compensatory programs that help tumor cells tolerate inflammatory pressure, oxidative stress, or therapy-associated damage(32, 35, 48, 56). The presence of GDF15 among the strongest IMRES-associated genes is particularly notable because tumor-derived GDF15 has been reported to limit LFA-1-dependent T-cell recruitment into tumors, providing one possible mechanism by which inflammatory tumors may still evade effective immune clearance(57, 58). The Hot+IMRES_high_ state further refined the meaning of immune-hot tumor-enriched epithelial regions. By integrating Composite Hotness with IMRES, Hot cells were separated into Hot+IMRES_high_ and Hot+IMRES_low_ populations, consistent with single-cell ecosystem and pathway-scoring frameworks used to resolve biologically distinct tumor states(29, 35, 45). The Hot+IMRES_high_ compartment represented the major immune-inflamed ecosystem subset and showed stronger checkpoint score, IFN/antigen-presentation signaling, redox adaptation, ferroptosis-associated transcript score, and radiation-stress signatures than Cold+IMRES_low_ cells(41, 49–51). The spatial permutation testing showed that Hot+IMRES_high_ cells were positioned closer to immune populations than expected under randomized labeling, supporting the idea that this state is spatially organized rather than randomly distributed(46, 52, 54). This does not prove direct immune engagement, tumor-cell killing, or causal signaling, but it does suggest that Hot+IMRES_high_ cells occupy tissue niches where immune proximity, checkpoint biology, and stress adaptation converge(7, 41, 47, 56).

Furthermore, the neighborhood analysis supports the idea that Hot+IMRES_high_ is a multicellular tissue state rather than a tumor-cell-only phenotype. Hot tumor-enriched epithelial regions were surrounded by immune, epithelial, and non-tumor neighboring compartments that also showed inflammatory and stress-associated features. This pattern suggests that Hot+IMRES_high_ is not simply a marker of immune infiltration, instead, it reflects a local ecosystem in which tumor-enriched epithelial cells and nearby cells share coordinated immune, metabolic, redox, and stress-response programs (6, 8, 12, 35, 46). This interpretation aligns with emerging spatial-omics studies showing that immune resistance is shaped by the joint organization of tumor-cell programs and immune-cell neighborhoods, rather than by immune-cell abundance alone. In this view, tumor-enriched epithelial states may help structure local immune architecture through antigen-presentation programs, checkpoint-associated signaling, metabolic remodeling, and stress-adaptive pathways. The rank-based analysis further strengthens this interpretation by serving as an internal robustness test for IMRES (43, 53, 59, 60). Benchmarking also showed that Composite Hotness, IMRES, IFN/antigen-presentation signaling, T-cell inflammation, rank-based IMRES, and CD274 expression could identify Hot+IMRES_high_ cells with meaningful discriminatory performance. Importantly, both IMRES and rank-based IMRES remained positively associated with immune and stress-response genes, including NFE2L2, GDF15, CD274, HLA-DRA, CXCL10, GPX2, TXNRD1, and KEAP1 (41, 43, 49, 51, 56, 58). Because rank-based methods rely on within-cell gene rankings and are less dependent on expression scale or normalization, this agreement supports the interpretation that the Hot+IMRES_high_ ecosystem is not solely an artifact of one scoring method (43, 44).

However, this should still be interpreted cautiously. Because all analyses were performed using one published Xenium resource, the rank-based analysis provides internal analytical support, not external validation. Independent HNSCC cohorts, spatial protein or metabolic assays, and outcome-linked datasets will be needed to determine whether Hot+IMRES_high_ regions are reproducible therapeutic resistance niches (28, 41, 49, 50, 56). Overall, these findings refine the meaning of an immune-hot state in HNSCC. An inflamed tumor region may still contain checkpoint-associated transcripts, oxidative-stress adaptation, and other stress-response programs that limit effective immune clearance. This coexistence may help explain why tumors with immune infiltration do not always respond to immune-mediated attack or therapy (5, 6, 28, 56). Thus, this extended study of prior spatial work highlights the development of a tumor-centered immune-metabolic-redox ecosystem model, in which immune activation, antigen presentation, checkpoint regulation, redox adaptation, ferroptosis-associated programs, and metabolic remodeling are organized together within local tumor neighborhoods.

Several limitations should be noted. The pathway scores were derived from a targeted Xenium transcript panel(38) and should be interpreted as transcript-level signatures, not direct measures of protein abundance, metabolite levels, metabolic flux, oxidative stress, mitochondrial function, NRF2 activation, ferroptosis, or radiation-stress signature. EpCAM^+^ enriched for tumor-associated epithelial compartments but did not establish malignant-cell identity without copy-number, mutation, or histopathologic validation(39, 40). Spatial proximity also indicates local association, not direct cell–cell contact, immune synapse formation, cytotoxic function, or causal communication. Finally, because Hot+IMRES_high_ was defined computationally from a public dataset, this state will need validation in independent HNSCC cohorts using orthogonal spatial, proteomic, metabolic, and outcome-linked approaches (35, 46), (59, 60).

## 5. Summary

Overall, this secondary analysis suggests that immune-hot HNSCC regions are not simply inflammatory niches, but spatial ecosystems where immune, metabolic, and redox programs overlap. The original McCord et al.(23) study established a spatial T-cell framework for HNSCC; the present study extends that resource by identifying a complementary epithelial ecosystem state in which immune activation, antigen presentation, checkpoint signaling, metabolic remodeling, and stress adaptation converge. IMRES provides a transcript-derived framework to quantify this biology, and the Hot+IMRES_high_ state highlights tumor neighborhoods where inflammatory and stress-adaptive programs coexist. This framework offers a focused basis for future studies of immune resistance, redox adaptation, and combination-therapy vulnerabilities in HNSCC.

## Data availability

This study was performed as a secondary analysis of an already published, publicly available spatial transcriptomic dataset generated by McCord et al. in 2026(23). The original Xenium dataset is available through the NCBI Gene Expression Omnibus under accession GSE300147. The GEO record includes 18 Xenium samples, consisting of 17 HNSCC sections and one ameloblastoma section, together with sample-level accessions and associated metadata. The processed tabular outputs generated during the current secondary analysis, including CSV summary tables and analysis outputs used to support the figures and results, are available from the corresponding author upon reasonable request.

## Ethics statement

This study used only publicly available, previously generated data. No new human specimens were collected, no new sequencing was performed, and no additional intervention involving human participants was conducted as part of this secondary analysis. Ethical approval, informed consent, and specimen acquisition for the original dataset are described in the original McCord et al. study(23).

## Code availability

Custom Python scripts used for Composite Hotness scoring, pathway scoring, metabolic ecosystem analysis, neighborhood analysis, IMRES generation, spatial permutation testing, rank-based robustness analysis, and benchmarking are available from the corresponding author upon reasonable request.

## Acknowledgements

The author acknowledges and thanks McCord et al.(23) for generating, curating, and publicly releasing the GSE300147 spatial transcriptomic dataset used in this study. The original work established a spatial T-cell receptor profiling framework in head and neck squamous cell carcinoma and created a valuable community resource that enabled subsequent secondary analyses. The present study is based entirely on reanalysis of the publicly available Xenium spatial transcriptomic data and would not have been possible without the efforts of the original investigators and study participants. The author also acknowledges the National Center for Biotechnology Information (NCBI) Gene Expression Omnibus for maintaining public access to the GSE300147 dataset and associated metadata.

## Author Contributions

K.S. conceived the study, performed data analysis, developed the computational workflow, interpreted the results, prepared the figures, and wrote and revised the manuscript.

## Disclosure for use of artificial intelligence (AI) tools

Microsoft Copilot 365 was used to assist with drafting and refining Python analysis scripts and for grammar, clarity, and language editing during manuscript preparation. All analyses, outputs, interpretations, and manuscript text were reviewed, verified, and approved by the author, who takes full responsibility for the accuracy and integrity of the work.

## References

1. Kirtane K, Dhar H, D’Cruz A, Yom SS. Emerging Treatments and Novel Modalities in Head and Neck Cancers. American Society of Clinical Oncology Educational Book. 2026;46(3):e516922.

2. Liu X, Harbison RA, Varvares MA, Puram SV, Peng G. Immunotherapeutic strategies in head and neck cancer: challenges and opportunities. J Clin Invest. 2025;135(8).

3. Puram SV, Rocco JW. Molecular Aspects of Head and Neck Cancer Therapy. Hematol Oncol Clin North Am. 2015;29(6):971–92.

4. Pan J, Chen S, Jin L, Chen K, Ye Z, Li B. Immunotherapy-driven remodeling of the tumor immune microenvironment: Spatiotemporal heterogeneity and multidimensional dynamics. Biochimica et Biophysica Acta (BBA) - Reviews on Cancer. 2026;1881(2):189544.

5. Blank C, Gajewski TF, Mackensen A. Interaction of PD-L1 on tumor cells with PD-1 on tumor-specific T cells as a mechanism of immune evasion: implications for tumor immunotherapy. Cancer Immunol Immunother. 2005;54(4):307–14.

6. Fu T, Dai LJ, Wu SY, Xiao Y, Ma D, Jiang YZ, et al. Spatial architecture of the immune microenvironment orchestrates tumor immunity and therapeutic response. J Hematol Oncol. 2021;14(1):98.

7. Li JR, Pan X, Lin Y, Zhao Y, Liu Y, Li Y, et al. Spatial Proximity of Immune Cell Pairs to Cancer Cells in the Tumor Microenvironment as Biomarkers for Patient Stratification. Cancers (Basel). 2025;17(14).

8. Blise KE, Sivagnanam S, Banik GL, Coussens LM, Goecks J. Single-cell spatial architectures associated with clinical outcome in head and neck squamous cell carcinoma. NPJ Precis Oncol. 2022;6(1):10.

9. Choi S-W, Kim JH, Hong J, Kwon M. Mapping immunotherapy potential: spatial transcriptomics in the unraveling of tumor-immune microenvironments in head and neck squamous cell carcinoma. Frontiers in Immunology. 2025;Volume 16 - 2025.

10. Steele K, Barnett C, Ladwa R, Donovan ML, Lawler C, Kulasinghe A. Tumour microenvironment diversity of HNSCC and the molecular landscape of recurrent disease. Oral Oncology. 2026;174:107864.

11. Jenkins E, Whitehead T, Fellermeyer M, Davis SJ, Sharma S. The current state and future of T-cell exhaustion research. Oxf Open Immunol. 2023;4(1):iqad006.

12. Cohn DE, Forder A, Marshall EA, Vucic EA, Stewart GL, Noureddine K, et al. Delineating spatial cell-cell interactions in the solid tumour microenvironment through the lens of highly multiplexed imaging. Front Immunol. 2023;14:1275890.

13. Qin X, Wu H, Pan J, Kang K, Shi Y, Bu S. Immune-metabolic crosstalk in HNSCC: mechanisms and therapeutic opportunities. Frontiers in Oncology. 2025;Volume 15 - 2025.

14. Zhang K-Y, Zhu X-Z, Yan Y-X, Ying X-H, Zhang J, Wang Z-H. Tumor evolution: signaling pathways, molecular mechanisms and therapeutic targets. Signal Transduction and Targeted Therapy. 2026;11(1):280.

15. Ma W, Wang Q, Guo L, Ju X. The molecular mechanisms, roles, and potential applications of PANoptosis in cancer treatment. Front Immunol. 2025;16:1550800.

16. Sharma B, Agnihotri N. Role of cholesterol homeostasis and its efflux pathways in cancer progression. The Journal of Steroid Biochemistry and Molecular Biology. 2019;191:105377.

17. Xie B, Guo Y. Molecular mechanism of cell ferroptosis and research progress in regulation of ferroptosis by noncoding RNAs in tumor cells. Cell Death Discov. 2021;7(1):101.

18. Shimura T. The role of mitochondrial oxidative stress and the tumor microenvironment in radiation-related cancer. Journal of Radiation Research. 2021;62(Supplement_1):i36–i43.

19. Su ZF, Wu CZ, Ding HR, Zhan Q, Ba YB, Cheng L, et al. Spatially resolved multiomics reveals the self-enforcing property of the leading-edge multicellular ecosystem of head and neck cancer. Proc Natl Acad Sci U S A. 2026;123(2):e2519474123.

20. Puram SV, Tirosh I, Parikh AS, Patel AP, Yizhak K, Gillespie S, et al. Single-Cell Transcriptomic Analysis of Primary and Metastatic Tumor Ecosystems in Head and Neck Cancer. Cell. 2017;171(7):1611–24.e24.

21. Mito I, Takahashi H, Kawabata-Iwakawa R, Ida S, Tada H, Chikamatsu K. Comprehensive analysis of immune cell enrichment in the tumor microenvironment of head and neck squamous cell carcinoma. Sci Rep. 2021;11(1):16134.

22. Hu Y, Liu W, Fang W, Dong Y, Zhang H, Luo Q. Tumor energy metabolism: implications for therapeutic targets. Mol Biomed. 2024;5(1):63.

23. McCord KA, Kan E, Hyslop S, Xia AY, Hofferek CJ, Lewis JS, Jr., et al. Single-cell TCR mapping reveals spatially coordinated T cell states in head and neck cancer. Sci Immunol. 2026;11(118):eaec3133.

24. Dong J, Xiao X, Nong L, Xue X, Wang L, Sun X, et al. Xenium-based spatial transcriptomic analyses uncover prognosis-associated heterogeneity in the tumor microenvironment (TME) of angioimmunoblastic T-cell lymphoma (AITL). J Pathol. 2026;269(1):71–85.

25. Luxenburger H, Thimme R, Hofmann M. T cell adaptation in chronic infections and tumors. Cellular & Molecular Immunology. 2026;23(5):440–56.

26. van der Leun AM, Thommen DS, Schumacher TN. CD8(+) T cell states in human cancer: insights from single-cell analysis. Nat Rev Cancer. 2020;20(4):218–32.

27. Lee HJ, Cho E, Song KH, Kim TW. Tumor Cells as Architects of Immune Refractoriness: Dismantling Intrinsic Programs of Tumor Cells for Clinical Translation. Immune Netw. 2026;26(1):e11.

28. Wang L, Geng H, Liu Y, Liu L, Chen Y, Wu F, et al. Hot and cold tumors: Immunological features and the therapeutic strategies. MedComm (2020). 2023;4(5):e343.

29. Wang J, Nagarajan P, Cho S, Liu Y, Seeley EH, Dai Y, et al. Integration of spatial single-cell proteomics and spatial metabolomics reveals tumor microenvironment predictive of immunotherapy response in mucosal melanoma. bioRxiv. 2026.

30. Liu G, Yao X, Hou Y, Deng W, Zhang L, Li S, et al. Metabolomic and transcriptomic profiling of HNSCC identifies AMIGO2 as a therapeutic target modulating tumor microenvironment. NPJ Precis Oncol. 2025;9(1):358.

31. Uhl B, Eggert D, Smiljanov B, von Thun Und Hohenstein K, Walz C, Kranz G, et al. Targeting EpCAM expression via near-infrared fluorescent antibodies enables microscopic delineation of primary and recurrent HNSCC. BMC Cancer. 2026;26(1).

32. Tang C, Xie AX, Liu EM, Kuo F, Kim M, DiNatale RG, et al. Immunometabolic coevolution defines unique microenvironmental niches in ccRCC. Cell Metab. 2023;35(8):1424–40.e5.

33. Wang K, Deng N, Tao Y, Yang X, Yuan M, Gao L, et al. Integrated single-cell and spatial transcriptomic analysis of T cell exhaustion and immunometabolic remodeling in HPV-positive oropharyngeal squamous cell carcinoma. bioRxiv. 2026.

34. Yuan Y. Spatial Heterogeneity in the Tumor Microenvironment. Cold Spring Harb Perspect Med. 2016;6(8).

35. Luca BA, Steen CB, Matusiak M, Azizi A, Varma S, Zhu C, et al. Atlas of clinically distinct cell states and ecosystems across human solid tumors. Cell. 2021;184(21):5482–96.e28.

36. Wolf FA, Angerer P, Theis FJ. SCANPY: large-scale single-cell gene expression data analysis. Genome Biol. 2018;19(1):15.

37. Virshup I, Bredikhin D, Heumos L, Palla G, Sturm G, Gayoso A, et al. The scverse project provides a computational ecosystem for single-cell omics data analysis. Nat Biotechnol. 2023;41(5):604–6.

38. Nerurkar SN, Goh D, Cheung CCL, Nga PQY, Lim JCT, Yeong JPS. Transcriptional Spatial Profiling of Cancer Tissues in the Era of Immunotherapy: The Potential and Promise. Cancers (Basel). 2020;12(9).

39. Lindgren G, Wennerberg J, Ekblad L. Cell line dependent expression of EpCAM influences the detection of circulating tumor cells with CellSearch. Laryngoscope Investig Otolaryngol. 2017;2(4):194–8.

40. Chikamatsu K, Takahashi H, Tada H, Uchida M, Ida S, Tomidokoro Y, et al. Circulating Tumor Cells in Head and Neck Squamous-Cell Carcinoma Exhibit Distinct Properties Based on Targeted Epithelial-Related Markers. Current Issues in Molecular Biology. 2025;47(4):240.

41. Ayers M, Lunceford J, Nebozhyn M, Murphy E, Loboda A, Kaufman DR, et al. IFN-γ-related mRNA profile predicts clinical response to PD-1 blockade. J Clin Invest. 2017;127(8):2930–40.

42. Allam M, Hu T, Lee J, Aldrich J, Badve SS, Gökmen-Polar Y, et al. Spatially variant immune infiltration scoring in human cancer tissues. NPJ Precis Oncol. 2022;6(1):60.

43. Aibar S, González-Blas CB, Moerman T, Huynh-Thu VA, Imrichova H, Hulselmans G, et al. SCENIC: single-cell regulatory network inference and clustering. Nature Methods. 2017;14(11):1083–6.

44. Wang Ruoqiao H, Thakar J. Comparative analysis of single-cell pathway scoring methods and a novel approach. NAR Genomics and Bioinformatics. 2024;6(3).

45. Xiao Z, Dai Z, Locasale JW. Metabolic landscape of the tumor microenvironment at single cell resolution. Nature Communications. 2019;10(1):3763.

46. Palla G, Spitzer H, Klein M, Fischer D, Schaar AC, Kuemmerle LB, et al. Squidpy: a scalable framework for spatial omics analysis. Nature Methods. 2022;19(2):171–8.

47. Lopez Janeiro A, Miraval Wong E, Jiménez-Sánchez D, Ortiz de Solorzano C, Lozano MD, Teijeira A, et al. Spatially resolved tissue imaging to analyze the tumor immune microenvironment: beyond cell-type densities. Journal for ImmunoTherapy of Cancer. 2024;12(5):e008589.

48. Pedrosa L, Foguet C, Oliveres H, Archilla I, de Herreros MG, Rodríguez A, et al. A novel gene signature unveils three distinct immune-metabolic rewiring patterns conserved across diverse tumor types and associated with outcomes. Front Immunol. 2022;13:926304.

49. Panieri E, Telkoparan-Akillilar P, Suzen S, Saso L. The NRF2/KEAP1 Axis in the Regulation of Tumor Metabolism: Mechanisms and Therapeutic Perspectives. Biomolecules. 2020;10(5).

50. Daolin T, Rui K. NFE2L2 and ferroptosis resistance in cancer therapy. Cancer Drug Resistance. 2024;7(0):41.

51. Yang K, Wang X, Song C, He Z, Wang R, Xu Y, et al. The role of lipid metabolic reprogramming in tumor microenvironment. Theranostics. 2023;13(6):1774–808.

52. Kumar G, Pandurengan RK, Parra ER, Kannan K, Haymaker C. Spatial modelling of the tumor microenvironment from multiplex immunofluorescence images: methods and applications. Frontiers in Immunology. 2023;Volume 14 - 2023.

53. Silkwood K, Dollinger E, Gervin J, Atwood S, Nie Q, Lander AD. Leveraging gene correlations in single cell transcriptomic data. BMC Bioinformatics. 2024;25(1):305.

54. Benimam MM, Meas-Yedid V, Mukherjee S, Frafjord A, Corthay A, Lagache T, et al. Statistical analysis of spatial patterns in tumor microenvironment images. Nature Communications. 2025;16(1):3090.

55. Pedregosa F, Varoquaux G, Gramfort A, Michel V, Thirion B, Grisel O, et al. Scikit-learn: Machine Learning in Python. Journal of Machine Learning Research. 2012;12.

56. Patel R, Saab K, Luo L, Ma Y, Osman RA, Williams NT, et al. Nrf2 Hyperactivation as a Driver of Radiotherapy Resistance and Suppressed Antitumor Immunity in Head and Neck Squamous Cell Carcinoma. Clin Cancer Res. 2025;31(19):4184–95.

57. Wischhusen J, Melero I, Fridman WH. Growth/Differentiation Factor-15 (GDF-15): From Biomarker to Novel Targetable Immune Checkpoint. Front Immunol. 2020;11:951.

58. Haake M, Haack B, Schäfer T, Harter PN, Mattavelli G, Eiring P, et al. Tumor-derived GDF-15 blocks LFA-1 dependent T cell recruitment and suppresses responses to anti-PD-1 treatment. Nature Communications. 2023;14(1):4253.

59. Minogue E, Baldominos P, Hsu L, Haigis MC, Agudo J. Decoding the spatial dynamics of tumor and immune cell interactions in solid cancers. Cancer Cell. 2026;44(1):1–5.

60. Liu Y, Dai Y, Wang L. Spatial omics at the forefront: emerging technologies, analytical innovations, and clinical applications. Cancer Cell. 2026;44(1):24–49.

