## Supplementary Table S1 for "Immune-metabolic-redox ecosystems define spatially organized tumor states in head and neck squamous cell carcinoma"

**Suppl Table 1.Primary IMRES Component Gene Sets**

| **Component** | **Candidate_genes** | **Genes_available_in_Xenium** | **Genes_missing_from_Xenium** | **N_candidate_genes** | **N_available_genes** | **Included_in_primary_IMRES** |
| --- | --- | --- | --- | --- | --- | --- |
| **T_Cell_Inflamed_Score** | CD3D;CD3E;CD4;CD8A;NKG7;PRF1;GZMB;CXCL9;CXCL10;CCL5 | CD3D;CD3E;CD4;CD8A;NKG7;PRF1;GZMB;CXCL9;CXCL10;CCL5 |  | 10 | 10 | TRUE |
| **Checkpoint_Score** | CD274;PDCD1;CTLA4;HAVCR2;TIGIT;LAG3;TNFRSF9;CD28;IL2RA | CD274;PDCD1;CTLA4;HAVCR2;TIGIT;LAG3;TNFRSF9;CD28;IL2RA |  | 9 | 9 | FALSE |
| **IFN_Antigen_Score** | STAT1;IRF1;HLA-DRA;HLA-DRB1;HLA-DPA1;HLA-DPB1;B2M;TAP1;CXCL9;CXCL10 | HLA-DRA;CXCL9;CXCL10 | STAT1;IRF1;HLA-DRB1;HLA-DPA1;HLA-DPB1;B2M;TAP1 | 10 | 3 | TRUE |
| **Lipid_Metabolism_Score** | LPL;APOA5;ADIPOQ;PPARG;PLIN4;HMGCS2;AKR1C3;PLA2G7;ABCC11 | LPL;APOA5;ADIPOQ;PPARG;PLIN4;HMGCS2;AKR1C3;PLA2G7;ABCC11 |  | 9 | 9 | TRUE |
| **Prostaglandin_Eicosanoid_Score** | PTGDS;HPGDS;PLA2G7 | PTGDS;HPGDS;PLA2G7 |  | 3 | 3 | TRUE |
| **Cholesterol_Sterol_Score** | HMGCS2;CYP1A1;CYP2A7;CYP2B6;CYP2F1;CYP3A4;CYP4B1 | HMGCS2;CYP1A1;CYP2A7;CYP2B6;CYP2F1;CYP3A4;CYP4B1 |  | 7 | 7 | TRUE |
| **Oxidative_Redox_Lipid_Stress_Score** | TXNRD1;GPX2;NFE2L2;KEAP1;GDF15 | TXNRD1;GPX2;NFE2L2;KEAP1;GDF15 |  | 5 | 5 | TRUE |
| **NRF2_Antioxidant_Score** | NFE2L2;KEAP1;GDF15;TXNRD1;GPX2 | NFE2L2;KEAP1;GDF15;TXNRD1;GPX2 |  | 5 | 5 | TRUE |
| **Ferroptosis_Score** | NFE2L2;KEAP1;GDF15;TXNRD1;GPX2 | NFE2L2;KEAP1;GDF15;TXNRD1;GPX2 |  | 5 | 5 | TRUE |
| **Mitochondrial_Stress_UPRmt_Score** | HSPD1;HSPE1;ATF5;DDIT3;HSPA9 |  | HSPD1;HSPE1;ATF5;DDIT3;HSPA9 | 5 | 0 | FALSE |
| Radiation_Stress_Damage_Score | GDF15;FAS;BCL2L11;CDKN1A;MDM2 | GDF15;FAS;BCL2L11;MDM2 | CDKN1A | 5 | 4 | TRUE |
| CD274_expr | CD274 | CD274 |  | 1 | 1 | TRUE |
