## Supplementary Table S2 for "Immune-metabolic-redox ecosystems define spatially organized tumor states in head and neck squamous cell carcinoma"

**Suppl Table 2. Rank IMRES Positive - Negative Gene List**

| **Gene** | **Spearman_r** | **Pvalue** | **Rank_IMRES_signature** |
| --- | --- | --- | --- |
| NFE2L2 | 0.507678233 | 0 | Rank_IMRES_Positive |
| GDF15 | 0.466344188 | 0 | Rank_IMRES_Positive |
| CYP4B1 | 0.419554764 | 0 | Rank_IMRES_Positive |
| HLA-DRA | 0.380584418 | 0 | Rank_IMRES_Positive |
| EHF | 0.377229174 | 0 | Rank_IMRES_Positive |
| CD274 | 0.361058863 | 0 | Rank_IMRES_Positive |
| AKR1C3 | 0.355725524 | 0 | Rank_IMRES_Positive |
| CLIC6 | 0.34702272 | 0 | Rank_IMRES_Positive |
| BCL6 | 0.329786413 | 0 | Rank_IMRES_Positive |
| PIGR | 0.322822914 | 0 | Rank_IMRES_Positive |
| SOX2 | 0.314518439 | 0 | Rank_IMRES_Positive |
| GPX2 | 0.305435418 | 0 | Rank_IMRES_Positive |
| TXNRD1 | 0.304093495 | 0 | Rank_IMRES_Positive |
| PPARG | 0.300810506 | 0 | Rank_IMRES_Positive |
| KEAP1 | 0.290412326 | 0 | Rank_IMRES_Positive |
| ERBB2 | 0.286179377 | 0 | Rank_IMRES_Positive |
| MDM2 | 0.278410831 | 0 | Rank_IMRES_Positive |
| TFPI | 0.277905996 | 0 | Rank_IMRES_Positive |
| ADAM28 | 0.269728278 | 0 | Rank_IMRES_Positive |
| CXCL10 | 0.266116411 | 0 | Rank_IMRES_Positive |
| SERPINB3 | 0.254893662 | 0 | Rank_IMRES_Positive |
| PTN | 0.254384186 | 0 | Rank_IMRES_Positive |
| CXCL9 | 0.251330284 | 0 | Rank_IMRES_Positive |
| MLPH | 0.250591487 | 0 | Rank_IMRES_Positive |
| TCIM | 0.247475087 | 0 | Rank_IMRES_Positive |
| SLC26A2 | 0.245490701 | 0 | Rank_IMRES_Positive |
| CFB | 0.242640475 | 0 | Rank_IMRES_Positive |
| BCL2L11 | 0.242115766 | 0 | Rank_IMRES_Positive |
| TMC5 | 0.237540236 | 0 | Rank_IMRES_Positive |
| FOXA1 | 0.217625787 | 0 | Rank_IMRES_Positive |
| LAMP3 | 0.21317828 | 0 | Rank_IMRES_Positive |
| CD8A | 0.208108458 | 0 | Rank_IMRES_Positive |
| TRAC | 0.203063544 | 0 | Rank_IMRES_Positive |
| CFHR3 | 0.199864917 | 0 | Rank_IMRES_Positive |
| CD3E | 0.199371792 | 0 | Rank_IMRES_Positive |
| TRBC1 | 0.195845538 | 0 | Rank_IMRES_Positive |
| CLCA2 | 0.188787756 | 0 | Rank_IMRES_Positive |
| TRBC2 | 0.186530368 | 0 | Rank_IMRES_Positive |
| GATM | 0.185510218 | 0 | Rank_IMRES_Positive |
| CD3D | 0.179088081 | 0 | Rank_IMRES_Positive |
| CCL5 | 0.178603263 | 0 | Rank_IMRES_Positive |
| PLIN4 | 0.176784184 | 0 | Rank_IMRES_Positive |
| PTPRC | 0.176492564 | 0 | Rank_IMRES_Positive |
| ACE2 | 0.17269443 | 0 | Rank_IMRES_Positive |
| PLA2G7 | 0.172276468 | 0 | Rank_IMRES_Positive |
| FAS | 0.165287983 | 0 | Rank_IMRES_Positive |
| CD2 | 0.162092476 | 0 | Rank_IMRES_Positive |
| CD4 | 0.158209243 | 0 | Rank_IMRES_Positive |
| TENT5C | 0.153022106 | 0 | Rank_IMRES_Positive |
| GZMA | 0.148392164 | 0 | Rank_IMRES_Positive |
| COL17A1 | -0.342939804 | 0 | Rank_IMRES_Negative |
| TNC | -0.240098379 | 0 | Rank_IMRES_Negative |
| CAVIN1 | -0.222505732 | 0 | Rank_IMRES_Negative |
| PDPN | -0.179983919 | 0 | Rank_IMRES_Negative |
| FHL2 | -0.164713376 | 0 | Rank_IMRES_Negative |
| MFAP5 | -0.137969104 | 0 | Rank_IMRES_Negative |
| APCDD1 | -0.1350533 | 0 | Rank_IMRES_Negative |
| EPCAM | -0.131686507 | ####### | Rank_IMRES_Negative |
| FSTL3 | -0.125861747 | ####### | Rank_IMRES_Negative |
| CAV1 | -0.120022748 | ####### | Rank_IMRES_Negative |
| HPV16_L1 | -0.116656716 | ####### | Rank_IMRES_Negative |
| GPC3 | -0.098256428 | ####### | Rank_IMRES_Negative |
| THBS2 | -0.094026927 | ####### | Rank_IMRES_Negative |
| MYLK | -0.086238931 | ####### | Rank_IMRES_Negative |
| HPV16_E6 | -0.084949638 | ####### | Rank_IMRES_Negative |
| GPRC5A | -0.081597871 | ####### | Rank_IMRES_Negative |
| DST | -0.080540945 | ####### | Rank_IMRES_Negative |
| FGFBP1 | -0.080153211 | ####### | Rank_IMRES_Negative |
| ADAMTS1 | -0.075495443 | ####### | Rank_IMRES_Negative |
| PVR | -0.073231499 | 1.49E-95 | Rank_IMRES_Negative |
| HPV16_E7 | -0.07118546 | 2.22E-90 | Rank_IMRES_Negative |
| MMRN2 | -0.069074959 | 3.36E-85 | Rank_IMRES_Negative |
| STC2 | -0.067468296 | 2.32E-81 | Rank_IMRES_Negative |
| KLK11 | -0.062082319 | 3.75E-69 | Rank_IMRES_Negative |
| CDH16 | -0.058504457 | 1.32E-61 | Rank_IMRES_Negative |
| CRISPLD2 | -0.053813687 | 2.18E-52 | Rank_IMRES_Negative |
| THY1 | -0.052611415 | 3.78E-50 | Rank_IMRES_Negative |
| HPV16_L2 | -0.051177253 | 1.52E-47 | Rank_IMRES_Negative |
| TNFRSF13B | -0.051164085 | 1.61E-47 | Rank_IMRES_Negative |
| LTBP2 | -0.051146203 | 1.73E-47 | Rank_IMRES_Negative |
| SERPINB2 | -0.049665501 | 7.08E-45 | Rank_IMRES_Negative |
| HPV16_E2 | -0.049572467 | 1.03E-44 | Rank_IMRES_Negative |
| ERG | -0.04742917 | 4.47E-41 | Rank_IMRES_Negative |
| PDGFRB | -0.047331393 | 6.50E-41 | Rank_IMRES_Negative |
| IL1R2 | -0.042964312 | 5.23E-34 | Rank_IMRES_Negative |
| GEM | -0.042698035 | 1.31E-33 | Rank_IMRES_Negative |
| VCAN | -0.042481111 | 2.77E-33 | Rank_IMRES_Negative |
| EDN1 | -0.039709748 | 2.72E-29 | Rank_IMRES_Negative |
| IL1RL1 | -0.038045054 | 5.06E-27 | Rank_IMRES_Negative |
| SNCA | -0.036722762 | 2.75E-25 | Rank_IMRES_Negative |
| EGFL7 | -0.032839736 | 1.53E-20 | Rank_IMRES_Negative |
| HPV16_E5 | -0.032206816 | 8.11E-20 | Rank_IMRES_Negative |
| SRPX | -0.03076049 | 3.25E-18 | Rank_IMRES_Negative |
| CNN1 | -0.03052077 | 5.90E-18 | Rank_IMRES_Negative |
| SNAI1 | -0.029786158 | 3.56E-17 | Rank_IMRES_Negative |
| C6orf118 | -0.029176372 | 1.53E-16 | Rank_IMRES_Negative |
| SFRP2 | -0.028913591 | 2.85E-16 | Rank_IMRES_Negative |
| RAPGEF3 | -0.028888469 | 3.02E-16 | Rank_IMRES_Negative |
| ANPEP | -0.028880748 | 3.08E-16 | Rank_IMRES_Negative |
| 240103_CAINNARLMF_TRA | -0.028689341 | 4.81E-16 | Rank_IMRES_Negative |
